# Mechanical competition promotes selective listening to receptor inputs to resolve directional dilemmas in neutrophil migration

**DOI:** 10.1101/2022.02.21.481331

**Authors:** Amalia Hadjitheodorou, George R. R. Bell, Felix Ellett, Daniel Irimia, Robert Tibshirani, Sean R. Collins, Julie A. Theriot

## Abstract

As neutrophils navigate complex environments to reach sites of infection, they may encounter obstacles that force them to split their front into multiple leading edges, raising the question of how the cell selects which front to maintain and which front(s) to abandon. Here we challenge chemotaxing HL60 neutrophil-like cells with symmetric bifurcating microfluidic channels, enabling us to probe the cell-intrinsic properties of their decision-making process. Using supervised statistical learning, we demonstrate that cells commit to one leading edge late in the decision-making process, rather than amplifying early pre-existing asymmetries. Furthermore, we use optogenetic tools to show that receptor inputs only bias the decision similarly late, once mechanical stretching begins to weaken each front. Finally, optogenetic attempts to reverse cell decisions reveal that, once an edge begins retracting, it commits to this fate, with the kinase ROCK limiting its sensitivity to inputs until the retraction is complete. Collectively our results suggest a “selective listening” model in which both actively protruding cell fronts and actively retracting cell rears have strong commitment to their current migratory program.

## INTRODUCTION

During immune surveillance, neutrophils use chemotaxis to navigate through highly complex tissue environments to reach sites of infection and inflammation. This process involves integrating directional cues detected by G-protein coupled receptors (GPCRs), as well as physical constraints of the environment, into an underlying autonomous polarity circuitry (Prentice-Mott et al., 2013; Weiner et al., 2007; Xu et al., 2003). Upon encountering obstacles, neutrophils may be forced to split their single leading edge into multiple competing fronts. Eventually cells select one front to maintain, and abandon the rest, in order to maintain their physical integrity and continue their movement.

Chemoattractants, such as N-formyl-methionyl-leucyl-phenylalanine (fMLF), activate signaling via the heterotrimeric GTPase Giα, which orients and reinforces morphologically and chemically distinct cell front and rear domains, leading to directed migration (Xu et al., 2003). Rac and Cdc42 are well-established cell front polarity regulators, inducing branched actin polymerization at the leading edge via activation of the Arp2/3 complex (Etienne-Manneville and Hall, 2002). At the cell rear, RhoA activity activates the cascade of ROCK and phosphorylation of myosin regulatory light chain to facilitate assembly of active myosin II bipolar filaments and actomyosin contraction (Xu et al., 2003). Locally, the cell front and rear domains are self-reinforcing via positive feedback, and at the same time are mutually antagonistic (Hind et al., 2016; Wang et al., 2002, 2013; Weiner et al., 2002; Xu et al., 2003).

This combination of positive and negative feedback effectively serves to enhance asymmetric cell polarization and polarity maintenance, promoting the formation of a single protruding front domain and one actively retracting rear domain (Jilkine et al., 2007; Marée et al., 2012). Mechanical coupling, through membrane tension, has been demonstrated to promote the de novo protrusion of only a single leading edge at a time, minimizing the likelihood of a cell spontaneously generating multiple cell fronts (Houk et al., 2012). However, it is not clear whether this same mechanism might also influence the resolution of multiple fronts formed by splitting, as seen when cells encounter obstacles.

Directional choices are the result of both cell-intrinsic and cell-extrinsic factors. Recently, microfluidic approaches have been leveraged to probe the role of different kinds of external asymmetries (Ellett et al., 2022; Moreau et al., 2019; Prentice-Mott et al., 2013; Renkawitz et al., 2019). For example, neutrophils usually choose the path with the steepest chemoattractant gradient (Ambravaneswaran et al., 2010; Ellett et al., 2022), and multiple types of leukocytes prefer larger sized channels (Renkawitz et al., 2019). In addition, neutrophils and immature dendritic cells preferentially select paths of least hydraulic resistance (*i.e.,* cells barotax), while *Dictyostelium* cells prioritize chemical rather than barotactic guidance cues (Belotti et al., 2020).

In the present work we designed symmetric channels, intentionally eliminating extrinsic biases, with the goal of identifying the cell-intrinsic factors that allow the cell to break symmetry when two equivalent fronts are forced to compete. In other words, how do cells resolve the competing influence of two distinct fronts if there is no external difference biasing their directional choice? Directional decision-making is an intrinsically noisy process, with considerable cell-to-cell variability. To overcome this inherent challenge, we standardized decision-making by challenging single HL60 neutrophil-like cells with a simplified oval-shaped obstacle, positioned symmetrically inside microfluidic channels (Fig. 1A). In this design cells make choices, not because of external asymmetry, as the two competing fronts encounter identical external environments, but because of the cells’ intrinsic ability to maintain their structural integrity.

**Figure 1:**
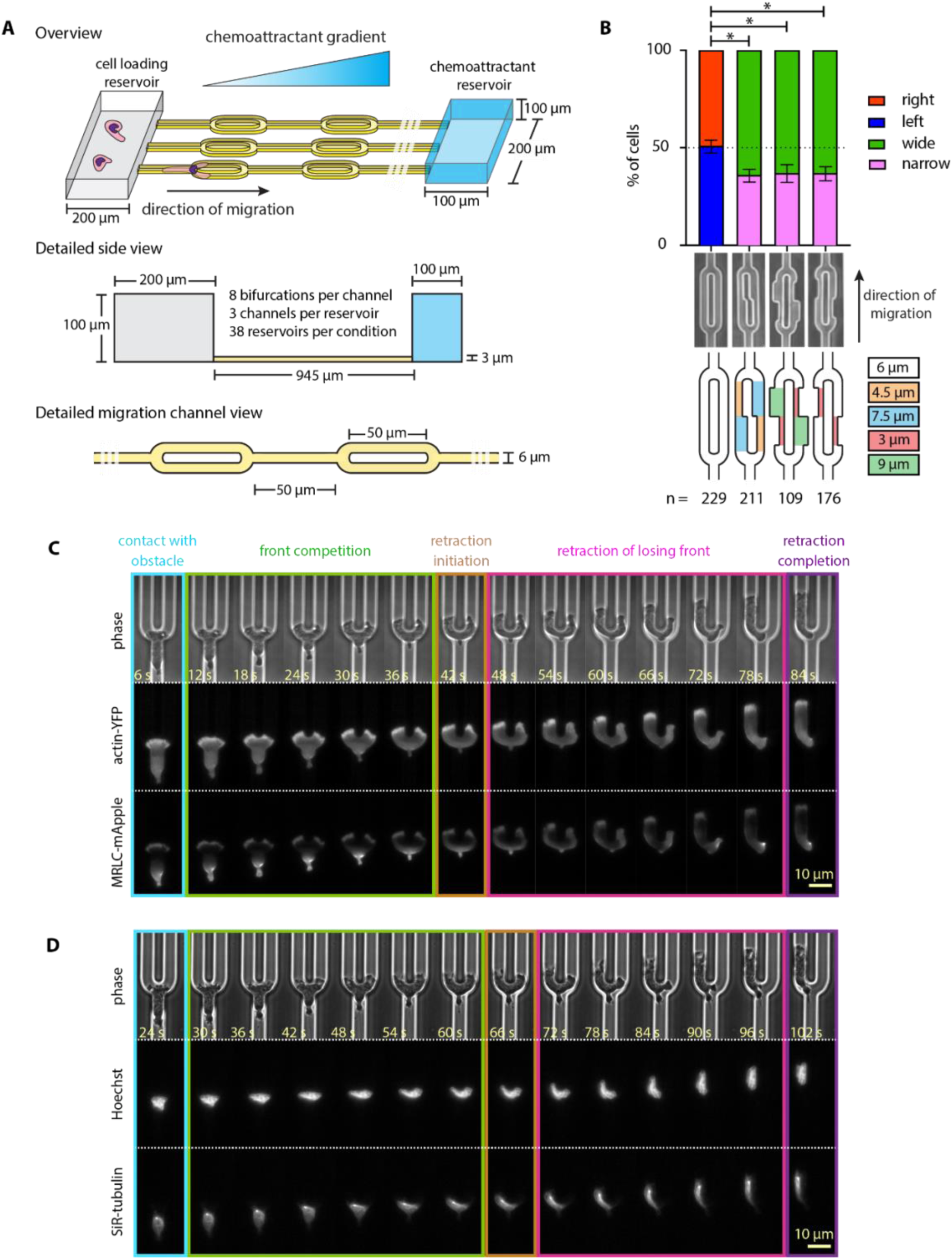
Cellular decision-making in an engineered symmetric environment. **(A)** Schematic illustration of the microfluidic device engineered to probe symmetric decision- making of HL60 cells. **(B)** Stacked bar plots of the percentage of cells that turned left vs. right (symmetric design) or wide vs. narrow (for all other channel designs). Error bars indicate standard deviations assuming a binomial distribution around the cumulative mean of each design. Two-sided Fisher exact tests revealed significant differences between the symmetric design and the asymmetric designs (* p < 0.05). **(C-D)** Representative live-cell imaging snapshots of single HL60 cells expressing actin-YFP and MRLC-mApple **(C)** and of the cell nucleus (stained with Hoechst) and microtubules (stained with SiR-tubulin) **(D)** during symmetric decision-making (Supplementary Movies 1-2). Images were captured every 3 s and subsampled for illustration purposes. Scale bar: 10 μm.

To determine whether pre-existing asymmetries in cellular structure might bias or predict the directional choice, and (if so) when during the process these asymmetries might arise, we leveraged a supervised statistical learning approach. Surprisingly, we found that we can predict the cell’s turning direction only late in the decision-making process, and we found no evidence that pre-existing asymmetries are amplified to help the cells decide which way to turn in this setting.

To further probe the cell’s commitment to protrusion and retraction programs, we used subcellular optogenetic receptor activation to locally activate Giα signaling (Bell et al., 2021) to bias the cell’s turning direction or reverse it. We found that stimulating a competing front only during the early phase of competition is insufficient to bias the cell’s direction, as each competing front appears fully committed to an independently self-reinforcing front protrusion program. As the cell stretches, the two fronts “weaken”, gradually decreasing their Cdc42 activity. During this later competition phase, optogenetic stimulation is sufficient to bias the cell direction. In addition, we found that, once a cell has made a directional choice, the losing front may enter a refractory period that requires complete retraction before stimulation can encourage a new protrusion; this refractory period is particularly pronounced for faster-moving cells. We show that the refractory nature of the retracting edge can be dampened by pharmacologically reducing the phosphorylation of myosin regulatory light chain and thus myosin II contractile activity. Collectively, our data suggest a context-dependent “listening” to receptor inputs, whereby both active cell fronts and cell rears have strong commitment to their current fate.

## RESULTS

### Cellular decision-making in an engineered symmetric environment

First, we sought to engineer a symmetric, tightly controlled and highly reproducible directional decision-making assay for HL60 cells. Our microfluidic device consists of an array of migration channels that connect a shared loading reservoir and chemoattractant chambers (Fig. 1A). The chemoattractant gradient is established along the migration channels via diffusion (see Methods). Cells enter the migration channels following a 100 nM fMLF gradient. The migration channels are 6 µm in width and 3 µm in height, forcing HL60 cells to occupy the entire cross-section, so that cells are mechanically confined and able to move only along the axis of the channels. Symmetric oval-shaped bifurcations are positioned every 50 µm (Fig. 1A). We found that cells presented with this symmetric bifurcation turned with equal probability to the left and to the right (Fig. 1B).

Cells responded to contact with the obstacle by splitting their front into two, with one active protrusion extending into each branch of the bifurcation. These two fronts competed, and the competition phase was resolved when one front started retracting, allowing the other side to “win” and guide the cell around the obstacle.

To gain better insight into how cell behavior depends on their mechanical constraints, we sought to quantify how directional bias changes as a function of channel geometry. To that end, we designed channels that have one wider and one narrower branch either immediately at the obstacle apex (2nd design from the left Fig. 1B) or at a 4 µm distance past the apex (3rd and 4rth designs from the left Fig. 1B). In designing channel geometries (Fig. 1B), we matched the hydraulic resistance between the two bifurcating branches, and only considered events where migrating cells did not have any other neighboring cells (preceding or following them) in the same channel. Cells showed a fairly modest (∼ 63%) but consistent preference for the initially wider branch, regardless of whether the asymmetry was introduced in the immediate vicinity of the obstacle apex or further away, suggesting that decisions are not finalized until cells have extended several micrometers into each channel. The preference of HL60 cells for wider channel branches is reminiscent of that of dendritic cells for larger sized channels (Renkawitz et al., 2019).

To investigate the cell-intrinsic factors giving rise to decision-making, we returned to the symmetric case and used fluorescent reporters for primary cytoskeletal and structural features to determine if their asymmetry might drive decision-making. Specifically, we used a neutrophil-like HL60 cell line stably expressing both actin-YFP and myosin regulatory light chain-mApple (abbreviated as MRLC-mApple hereafter) and observed actin and myosin localization dynamics during decision-making (Supplementary Movie 1). As expected, actin is enriched at both of the two competing fronts, while MRLC strongly localizes at the original cell rear. Accumulation of myosin at the losing front follows after the decision has been made (Tsai et al., 2019) (Fig. 1C). Labeling the nucleus with Hoechst and the microtubules with SiR-tubulin, we saw that in most events the nucleus and the microtubule organizing center are localized behind the obstacle apex at the moment of retraction, a behavior that is quite distinct from that exhibited by dendritic cells inside sharp Y-shaped obstacles (Renkawitz et al., 2019) (Fig. 1D, Supplementary Movie 2). Visual inspection revealed no obvious pre-existing asymmetries in cell shape, nucleus positioning, or in the spatial distributions of actin, myosin or microtubules that could reliably predict the directional choice.

### Faced with a symmetric dilemma, cell asymmetries arise only late in the decision-making process

To determine whether small asymmetries in cellular structure might bias or predict the decision-making, and (if so) when during the process these asymmetries might arise, we designed an agnostic statistical learning approach. Briefly, we collected time-lapse fluorescence microscopy movies of hundreds of decision-making events from single cells, capturing the organization and dynamics of different cytoskeletal components as well as cell shape-related information (Fig. 2A). We collected two datasets: one dataset composed of phase contrast images, along with fluorescence images of actin-YFP and MRLC-mApple, and another dataset composed of phase contrast images together with fluorescence images of the cell nucleus (Hoechst) and microtubules (SiR-tubulin). For both datasets, we collected over 160 decision-making events from single cells. For each decision-making event, we pre-processed the raw images and then proceeded to extract hundreds of features for each image using the open-source software CellProfiler (Carpenter et al., 2006; McQuin et al., 2018). These features report on shape and fluorescence intensity distribution metrics, as well as image texture and granularity.

**Figure 2:**
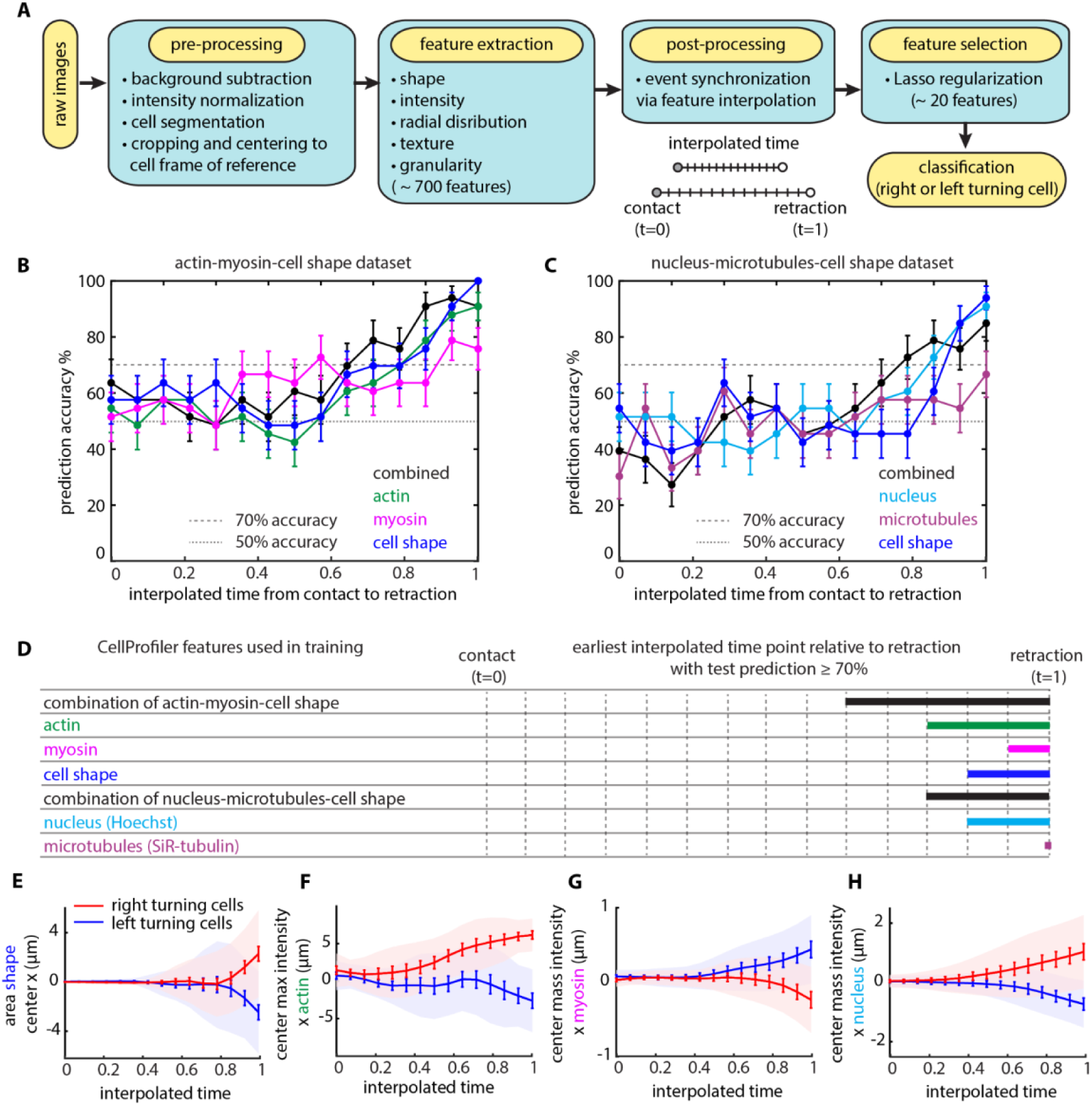
Faced with a symmetric dilemma, cell asymmetries arise only late in the decision- making process. **(A)** Flow diagram of the statistical learning pipeline designed to determine whether small asymmetries in cellular structure bias or predict the turning direction of HL60 cells during symmetric decision-making. **(B-C)** Test set prediction accuracy of statistical models leveraging information contained in binary masks and fluorescent images of actin and myosin **(B)** and in binary images and fluorescent images of the cell nucleus and microtubules **(C)** across interpolated time points between contact with the obstacle and retraction initiation. **(D)** Graphic representation of the earliest time point, relative to retraction initiation, with test prediction accuracy ≥ 70% for each statistical model. **(E-H)** Representative features with the most significant lasso predictive capacity; cell center *x*-coordinate **(E)**, maximum intensity of actin **(F)**, center of mass intensity (*i.e.,* intensity weighted centroid) of myosin **(G)** and of cell nucleus **(H)** over interpolated time between contact with the obstacle and retraction initiation.

We found that different cells make their decisions at different rates (Supplementary Fig. 1A), with decision-making time showing a negative correlation with migration speed before contact with the obstacle (Supplementary Fig. 1B). That is, cells that were moving quickly before contact with the obstacle also typically made their decision faster than did slower-moving cells. To facilitate comparisons across the large population of individual cells, we filtered the data to include only events whose time from contact with the obstacle until retraction initiation was within one standard deviation of the population average. This resulted in a final dataset size with 100 events in each dataset. To synchronize the filtered decision-making events, we worked on a rescaled time, interpolating feature values for a fixed number of equally spaced time points between contact with the obstacle and retraction initiation (Fig. 2A; see Methods).

For each dataset, out of the 100 decision-making events we randomly selected 67 for training and held back the remaining 33 as an independent test set. Logistic regression with “least absolute shrinkage and selection operator” (lasso) regularization was enforced to prevent overfitting the data (Tibshirani, 1996). Briefly, lasso regularization applies a penalty to the feature magnitudes to enforce use of only a limited number of nonzero features, leading to a sparse solution (see Methods). Leveraging lasso, we kept ∼ 10 features for training our binary (left/right) classification (Fig. 2A).

Plotting the prediction accuracy for the independent test set for each dataset, we see that at the time the cell contacts the obstacle (t=0) no statistical learning model predicts the cells’ ultimate direction with accuracy better than 50% (random) (Figs. 2B-2C). We also separately interrogated the cell shape, actin, myosin, nucleus and microtubule features, but found that models constructed on individual feature sets have poorer predictive power than models constructed by combining features from different cell structures (Figs. 2B-2D). Overall, the statistical model that we derived from the combination of cell shape, actin, and myosin features was the most predictive of the cell’s final turning direction. However, even with that best model, we were surprised to find that we can predict the cell’s turning direction with accuracy greater than 70% only during the last third of the decision-making process (typically about 15 s before the initiation of retraction).

Notably, the statistical model constructed on the microtubule features alone was the least predictive out of the ones examined. Microtubules have been shown to be indispensable for dendritic cell decision-making (Kopf et al., 2020; Renkawitz et al., 2019). Nocodazole treatment, known to depolymerize microtubules, resulted in a prominent phenotypic change whereby dendritic cells were unable to initiate a retraction event, often resulting in cell breaking (Kopf et al., 2020; Renkawitz et al., 2019). However, when we treated HL60 cells with nocodazole we did not observe such a retraction initiation defect nor cell fragmentation (Supplementary Figs. 1E-1F, Supplementary Movie 3). These results, together with the lack of predictive information associated with microtubule-based image features, suggests that HL60 cells resolve directional dilemmas using a mechanism that is distinct from that of dendritic cells.

The most highly scored individual features, as selected from the statistical models, also appear to diverge from symmetry (y=0) late in the decision-making process (Figs. 2E-2H). Different regularization methods, *e.g.,* elastic net (Zou and Hastie, 2005) or sparse group lasso (Simon et al., 2013) (see Methods), did not yield models of improved prediction accuracy (Supplementary Figs. 1C-1D).

Our statistical analysis suggests that any pre-existing cell-intrinsic asymmetries, if they exist, are not reflected in the structural organization of the cytoskeleton (actin, myosin, microtubules) or the positioning of the nucleus until late in the decision-making process. This conclusion is also consistent with our earlier finding that cells can detect asymmetries in channels at least 4 µm past the obstacle apex just as efficiently as asymmetries proximal to the apex (Fig. 2B). Overall, we conclude that neutrophils resolve symmetric directional dilemmas by random choice late in the decision-making process, and we find no evidence that pre-existing asymmetries are amplified to help the cells decide which way to turn.

### Optogenetic GPCR activation can only bias decision-making late in the process

Since cells challenged with a symmetric decision-making task appear to decide late into the process, we wondered whether receptor inputs could create early asymmetry to bias and speed up decisions. We used an HL60 cell line stably expressing parapinopsin (a light-sensitive Giα- family GPCR) and a spectrally compatible tdTomato/tdKatushka2 FRET biosensor for Cdc42 activity (Bell et al., 2021), a critical signaling component at the cell front (Lämmermann et al., 2009; Srinivasan et al., 2003; Van Keymeulen et al., 2006; Yang et al., 2016). This system allows direct measurements of signaling responses in cells whose migration is guided by parapinopsin (Bell et al., 2021) (Fig. 3A). Cells were confined to migrate inside the same symmetrically bifurcating microfluidic channels described above. Using a 407 nm laser, we locally stimulated cells with 10 ms light pulses, and with spatial precision of about a 1 μm diameter spot, using real-time image analysis to position the activation spot automatically and dynamically at the appropriate location (see Methods).

**Figure 3:**
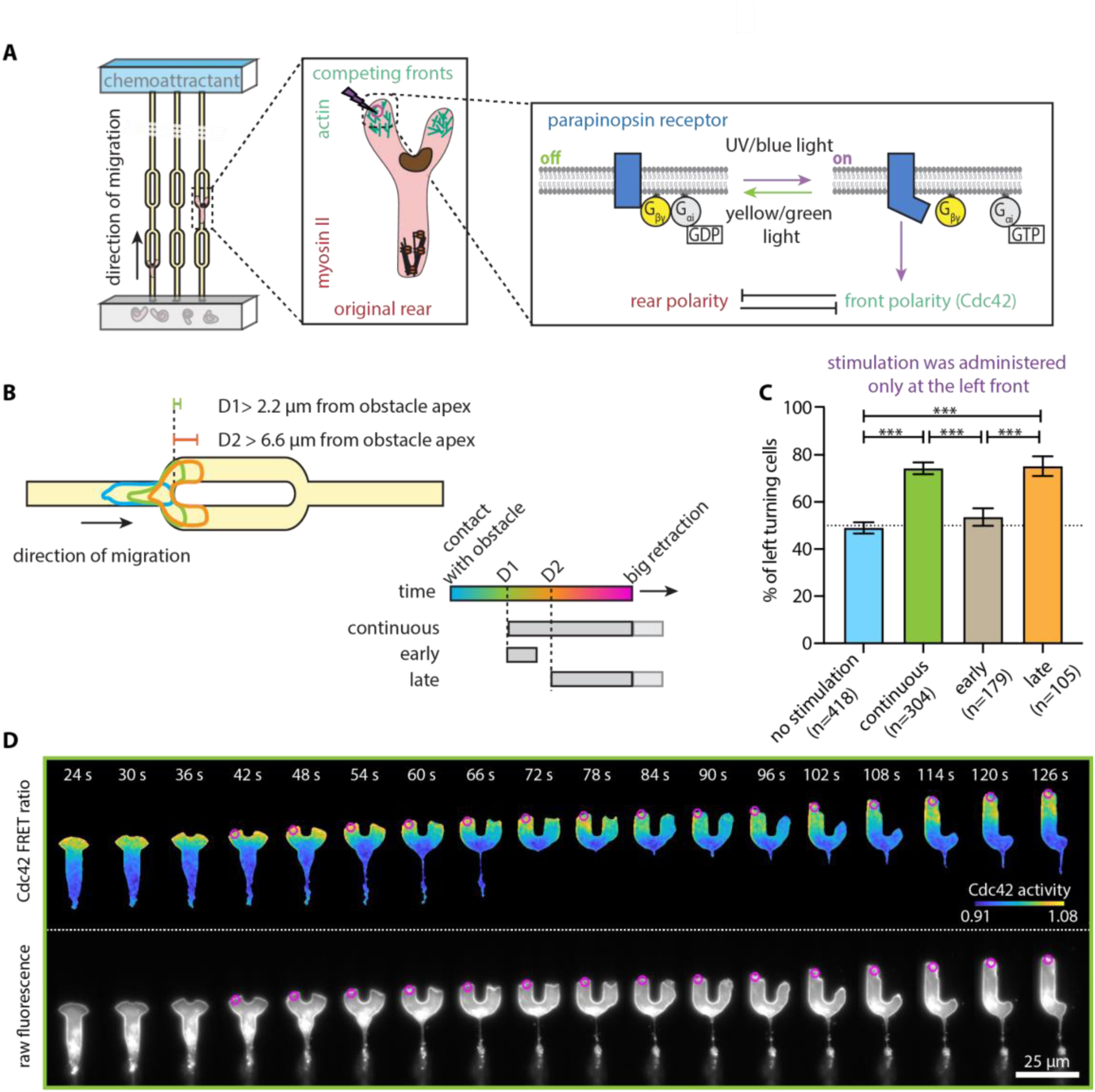
Optogenetic GPCR activation can only bias decision-making late in the process. **(A)** Schematic representation of optogenetic GPCR activation during the competition. HL60 cells expressing parapinopsin, an optogenetic receptor, migrate inside the symmetrically bifurcating microfluidic channels. The opsin is turned on using UV/blue light and off with yellow/green light. Blue light stimulation (magenta circle/lightning bolt) at the left competing cell front activates the receptor. The receptor is coupled to the polarity signal transduction network and activation reinforces the front polarity module. **(B)** Schematic of the competition stimulation assays. Stimulation is initiated after the two competing fronts have clearly separated from one another (*i.e.,* when both have extended at least 2.2 µm past the apex) for the continuous and early assays; late stimulation assay starts when both sides have extended at least 6.6 µm past the apex. Continuous and late stimulation entails persistent pulsated stimulation, whereas early stimulation comprises 5 pulses. **(C)** Bar plots of the percentage of left turning cells under no stimulation (n = 418 cells), continuous (n = 304 cells), early (n = 179 cells) and late (n = 105 cells) stimulation. Error bars indicate standard deviations assuming a binomial distribution around the cumulative mean of each assay. Two-sided Fisher exact tests revealed significant differences across no stimulation and continuous and late stimulation assays (*** p < 0.001). **(D)** Live-cell imaging snapshots of a cell expressing parapinopsin and a red/far-red Cdc42 FRET sensor. Cells migrate without experimental perturbation prior to initiation of continuous stimulation (magenta circles) at the left competing front (Supplementary Movie 4). Upper and lower panels show the registered sensors (gray scale) and the computed Cdc42 sensor FRET ratio, respectively. Images were captured every 3 s and subsampled for illustration purposes. Scale bar: 25 μm.

We found that repeatedly stimulating the left cell front every 3 s (beginning when both had extended at least 2.2 µm past the apex), was sufficient to bias the turning direction (75% left turning cells vs. 49% under no stimulation) (Figs. 3B-3C, Supplementary Movie 4). Given previous experimental results using optogenetic stimulation of this kind (Bell et al., 2021; Hadjitheodorou et al., 2021), we were pleased but not surprised that (this “continuous stimulation” assay was sufficient to bias the turning direction of a cell population. In addition, the continuous optogenetic GPCR activation resulted in marginally greater directional biasing as compared to the channel geometry asymmetries examined before (Fig. 1B).

However, we noticed that stimulation did not resolve the competition immediately (Fig. 3D). Conceptualizing the cell extending two fronts as a seesaw, perfectly balanced at the beginning of the competition, we wondered whether introducing a bias early in the process would be sufficient to tilt the seesaw so that the cell will follow that designated path in the future. To probe this question, we devised an “early stimulation” assay, delivering 5 stimulation pulses early on during the decision-making process, with the same stimulation onset as in the continuous experiment (Supplementary Movie 5). We discovered that this early intervention (in an attempt to bias signaling before the development of overt cytoskeletal structural asymmetries) had no effect, with the direction of the final behavioral decision returning to the stochastic 50%-50% left-right baseline. This result indicates that inducing a transient signaling asymmetry at an early stage is not sufficient to drive the system out of its balanced state.

Our results thus far suggested that decision-making happens late, and we reasoned that this may also be the time during which cells “listen” most carefully to receptor inputs. To test this directly, we devised a “late stimulation” assay (Supplementary Movie 6), where we stimulated the left front only after the competing fronts had extended 3-fold further than the continuous and early stimulation assays (about 6.6 µm past the apex of the obstacle) (Figs. 3B-3C). We found that late stimulation was just as effective as the continuous stimulation, providing additional support that early stimulation is futile and that optogenetic GPCR activation can bias decision-making only late in the process.

### Decision-making follows a stereotypical course dominated by autonomous polarity and cell stretching

Next, we sought to identify the underlying mechanism that allows the cells to become more amenable to directional bias late into their decision-making process. We began by determining whether changes in signaling activity precede and might determine the symmetric breaking in protrusion at the two fronts. We quantified the mean protrusion speed and magnitude of the Cdc42 activity of the two competing fronts for left-turning unstimulated control cells (Figs. 4A-4B). Examining the first half of the decision-making process (from 0 to 0.5 in interpolated time), we could detect no obvious difference either in speed or in Cdc42 activity between the two competing fronts. These observations are reminiscent of our analysis described above, where cytoskeletal asymmetries could be detected only very late in the decision-making process.

**Figure 4:**
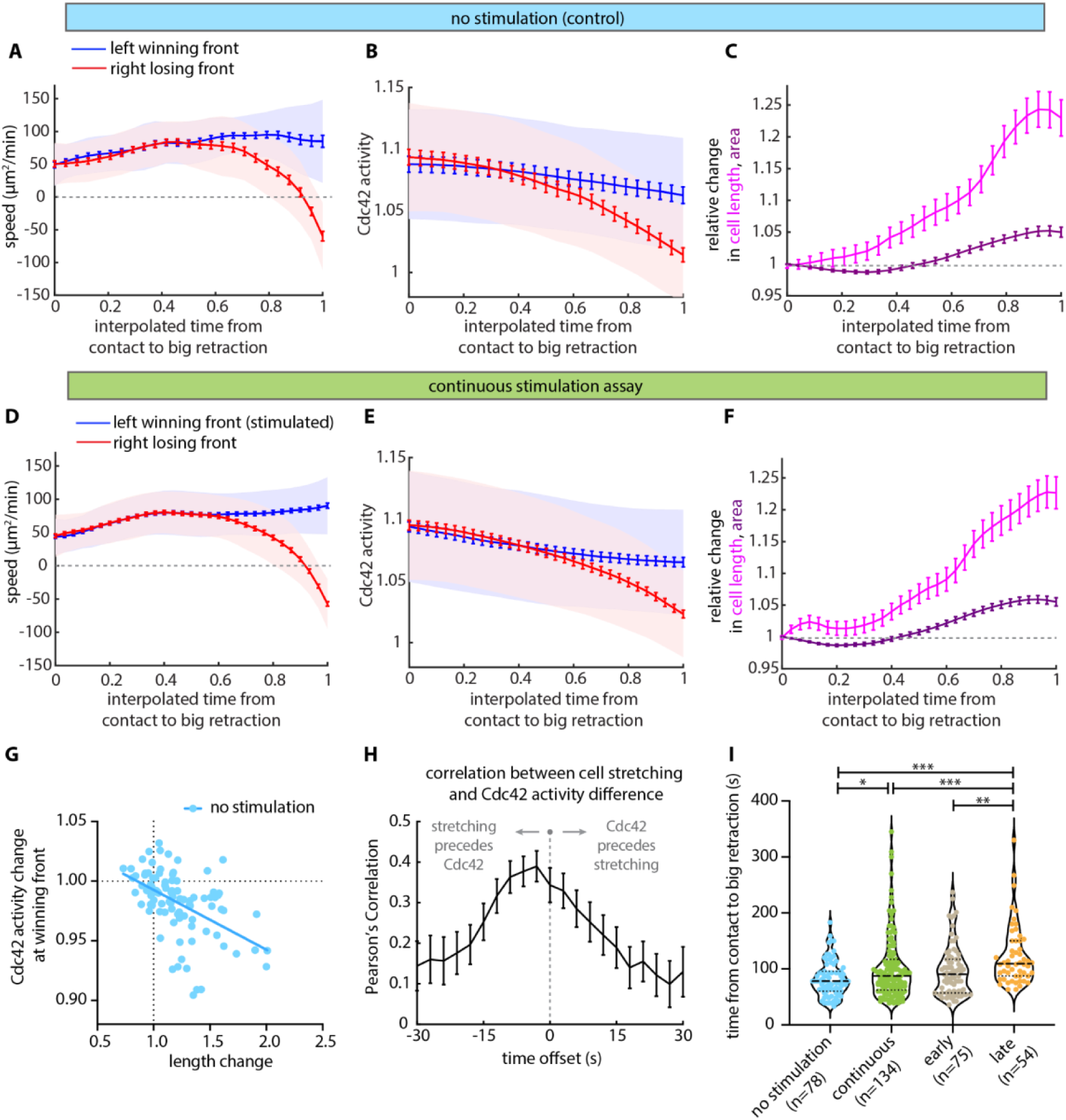
Decision-making follows a stereotypical course dominated by autonomous polarity and cell stretching. Mean speed **(A)** and magnitude of the Cdc42 activity (FRET ratio) **(B)** of the winning (blue) and the losing (red) competing fronts for 48 spontaneously left turning cells that received no stimulation (lines: means, shaded regions: SD, error bars: mean value ± SE). **(C)** Relative change in cell length (magenta) and cell area (purple), both normalized to the initial length and area at the time of contact with the obstacle, for 100 unstimulated (control) cells (lines: means, error bars: mean value ± SE). Mean speed **(D)** and magnitude of Cdc42 activity **(E)** of the winning (blue) and the losing (red) competing fronts for 125 stimulated left turning cells (lines: means, shaded regions: SD, error bars: mean value ± SE). **(F)** Relative change in cell length (magenta) and cell area (purple), both normalized to the initial length and area at the time of contact with the obstacle, for 166 continuously stimulated cells (lines: means, error bars: mean value ± SE). **(G)** Scatter plot of Cdc42 activity sensor fold-change and at the winning front vs. length change of 94 unstimulated cells. Changes were calculated from when the two competing fronts had clearly separated from one another (*i.e.* when both had extended at least 2.2 µm past the apex) and until big retraction initiation. Linear best-fit curve has been provided to guide the eye. **(H)** Temporal cross-correlation between the relative change in cell length and the absolute magnitude of the difference in Cdc42 activity between the winning and losing fronts as a function of time offset, for 75 unstimulated (control) cells (line: mean, error bars: mean value ± SE). The temporal analysis was performed for the same observation window as panel **(G)**. **(I)** Violin plots of the time from contact with the obstacle to big retraction of cells under no stimulation (n = 418 cells), continuous (n = 244 cells), early (n = 69 cells), and late (n = 97 cells) stimulation; *p*-values of two-sided Wilcoxon’s rank-sum test (\**p* < 0.05, \*\**p* < 0.01, \*\*\**p* < 0.001). For this analysis, cells were filtered so that all cells analyzed would have initiated the late stimulation protocol, if given the chance. In other words, only cells that extended their competing fronts at least 6.6 µm past the obstacle apex.

About 60% of the way through the process, both the speed and the Cdc42 activity of the two competing fronts began to diverge. Nevertheless, the losing front continued to move forward, just at a slower speed than the winning front, until the forward progress of the losing side stalled and began to retract about 90% of the way through the decision-making process (Fig. 4A). The deviation of Cdc42 signaling between the two competing fronts was similarly subtle and gradual (Fig. 4B). Notably, during this competition phase, the absolute amount of Cdc42 activity gradually decreased for both the left and right sides, but the rate of the activity decreased slightly more rapidly for the losing side (Fig. 4B). In addition, we observed gradual stretching of the cells during their decision-making (Fig. 4C).

Comparing left-turning cells whose left front had been stimulated (Figs. 4D-4F) with cells that spontaneously turned left with no stimulation (Figs. 4A-4C), we were surprised to find no measurable differences in cell stretching, protrusion speed, or the magnitude of the Cdc42 activity at either the winning or the losing front. These results are consistent with a model in which a strong self-organizing property of the polarity machinery drives the behavioral response. Therefore, even though continuous (and late) optogenetic GPCR activation clearly influences the turning direction across the population of cells, our results suggest that the autonomous polarity program dominates at the level of signaling in single cells, and that parapinopsin activation is only biasing the polarity machinery instead of completely overwriting it.

Even so, the fact that Cdc42 activity gradually declines at both fronts during the competition suggests that front polarity may be weakening. We hypothesized that mechanical cell stretching could be causing this weakening and thereby mediating the competition. To test this, we analyzed stretching and signaling at the single cell level. We found that the degree of cell stretching correlates with the drop in Cdc42 activity (Fig. 4G). Notably, cells that did not have apparent stretching showed very little change in Cdc42 activity, but cells that stretched to larger degrees tended to have correlated decreases in front signaling activity (Fig. 4G). Furthermore, cross-correlation analysis indicates that stretching is temporally correlated with and slightly precedes the divergence in Cdc42 signaling between the two fronts (Fig. 4H). Our results are consistent with previous findings that aspiration-mediated increase in mechanical tension in neutrophil-like cells is sufficient to cause long-range inhibition of actin assembly and Rac activity at the cell front (Houk et al., 2012), and they suggest that stretching may mediate competition by weakening both fronts and making them more amenable to external biasing.

Finally, when comparing the duration of the decision-making process, measured from contact with the obstacle to onset of big retraction (projected retraction area larger than 7.2 µm^2^), continuous and late stimulation assays show longer time averages, with more decision-making events requiring extended time to resolve, as compared to spontaneous resolution of the dilemma in unstimulated cells (Fig. 4I). That is, stimulation has a clear measurable effect on the system, but only by prolonging the decision-making process. Simply said, stimulation makes cells “think” longer before committing to a new migration direction.

### A committed retracting edge ignores input until it is fully retracted

Finally, we asked whether we could rescue a losing front and reprogram it to start protruding again. To that end, we let cells spontaneously pick a direction, but once we detected a big retraction (projected retraction area larger than 7.2 µm^2^), we started repetitively stimulating the retracting side. We called this assay “continuous stimulation” (Fig. 5A, Supplementary Movie 7). Similar to the competition assays described above, we also devised an “early stimulation” assay, where we stimulated the retracting edge with only 5 pulses (Supplementary Movie 8), as well as a “late stimulation” assay, where we let the cell retract all the way back to the apex of the obstacle and then began repetitively stimulating the previously retracting side (Fig. 5A, Supplementary Movie 9). We found that continuous stimulation was sufficient to reverse 27% of cells (Fig. 5B). Early stimulation was much less effective, only reversing 7% of cells, whereas late stimulation was even more effective than continuous stimulation, reversing 37% of cells (Fig. 5B).

**Figure 5:**
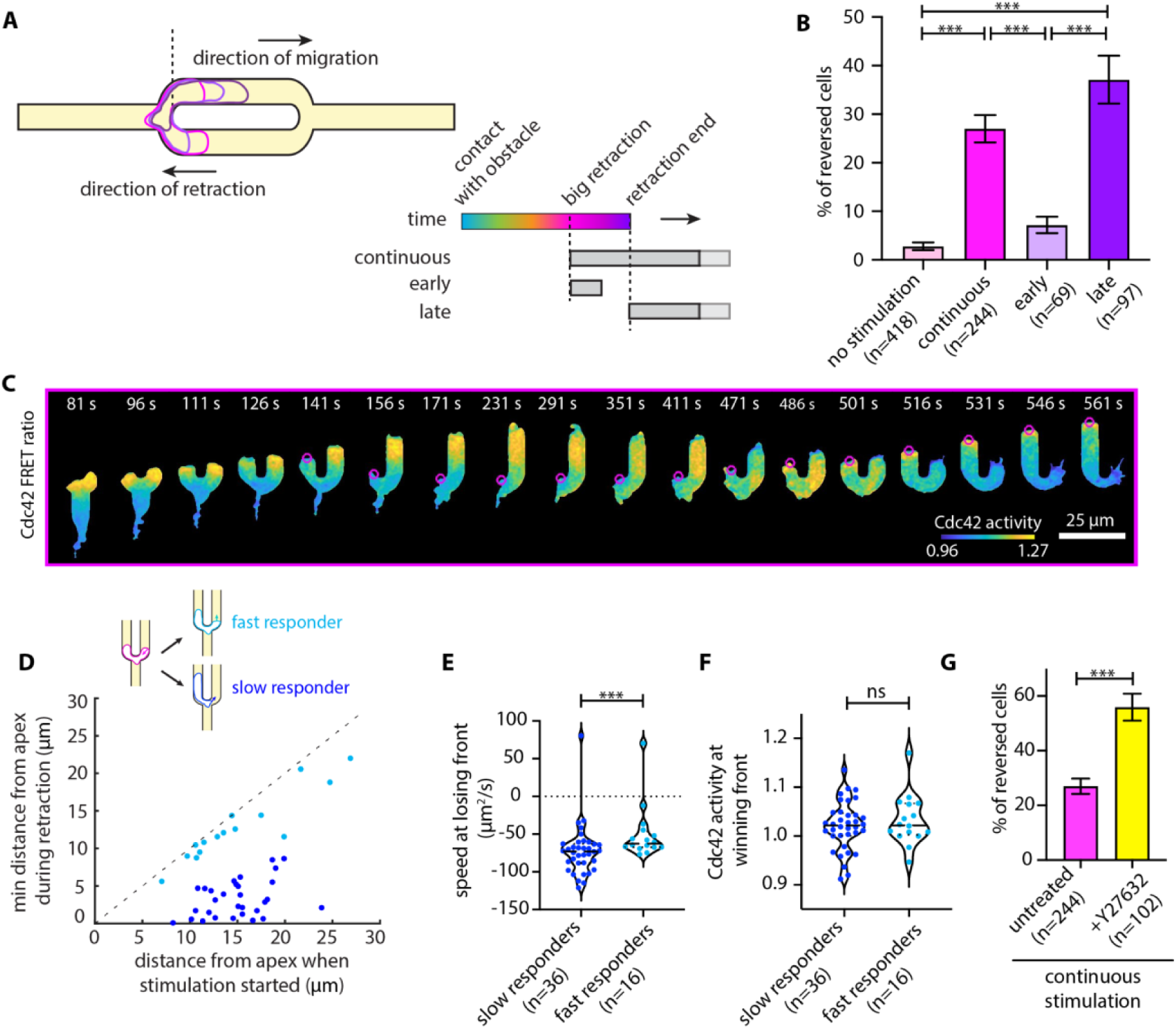
A committed retracting edge ignores input until it is fully retracted. **(A)** Schematic illustration of the reversal stimulation assays. Stimulation initiation commences after we detect a big retraction of a competing front (*i.e.,* retraction > 7.2 µm^2^) for the continuous and early assays; late stimulation assay commences when the retracting edge has reached the apex midline. Continuous and late stimulation entails persistent pulsated stimulation, whereas early stimulation comprises 5 pulses. **(B)** Bar plots showing the percentage of reversing cells under no stimulation (n = 418 cells), continuous (n = 244 cells), early (n = 69 cells) and late (n = 97 cells) stimulation. Error bars indicate standard deviations assuming a binomial distribution around the cumulative mean of each assay. Two-sided Fisher exact tests revealed significant differences across no stimulation and continuous and late stimulation assays (*** p < 0.001). **(C)** Live-cell imaging snapshots of a cell expressing parapinopsin and a red/far-red Cdc42 FRET sensor. Cells migrate and decide on a direction of turning without experimental perturbation. Upon identifying a big retraction, continuous stimulation (magenta circles) at the retracting/losing front commences (Supplementary Movie 7). Panels show the computed Cdc42 sensor activity. Images were captured every 3 s and subsampled for illustration purposes. Scale bar: 25 μm. **(D)** Scatter plot of minimum distance from the obstacle apex during retraction of the losing front vs. distance of the same front from the obstacle apex when the stimulation started, for 45 reversing cells. Cells stratified as slow (blue) and fast (cyan) responders. **(E-F)** Violin plots of speed of the losing/retracting front **(E)** and Cdc42 activity at the winning front **(F)** at the time of big retraction initiation for slow (n = 36 cells) and fast responders (n = 16 cells) under the continuous reversal assay; *p*-values of two-sided Wilcoxon’s rank-sum test (*** *p* < 0.001, ns: *p* > 0.05). **(G)** Bar plots of the percentage of reversing cells under no treatment (n = 244 cells) and Y2763 treatment (n = 102 cells). Error bars indicate standard deviations assuming a binomial distribution around the cumulative mean of each condition Two-sided Fisher exact tests revealed a significant difference between conditions (*** p < 0.001).

Even for the cells that were amenable to optogenetic reversal, we found that the response did not occur immediately. Specifically, we found that 71% of reversing cells, despite our continuous stimulation, kept retracting their losing front all the way back to the obstacle apex before they started reversing (Figs. 5C-5D). For these slow-responding cells, the losing front appears to enter a refractory period that requires complete retraction to the cell body before stimulation can reorient them. By classifying cells into slow and fast responders, we found that the slow responders typically had a faster retracting speed at the start of stimulation (Fig. 5E). We found no difference in the magnitude of Cdc42 activity at the winning front between slow and fast responders (Fig. 5F). That is, it appears that a losing front with high retraction speed tends to have a strong “conviction” to keep retracting, and only after it has completed its retraction program will it become amenable to new receptor inputs and switch into a protrusive behavior, ultimately leading to cell reversal.

Prior work has shown that myosin II contractile activity at the cell’s rear contributes to the maintenance of polarity, while also inhibiting cell front polarity programs (Hadjitheodorou et al., 2021; Smith et al., 2007; Uchida et al., 2003; Xu et al., 2003). We therefore sought to test the effects of altering myosin contractility in our optogenetic reversal assay. We found that ROCK inhibition with Y27632 treatment, known to decrease MRLC phosphorylation and myosin II activity in neutrophils (Niggli, 1999; Tsai et al., 2019), increases the percentage of reversing cells (Fig. 5G). Similarly to what we previously found for optogenetic receptor-directed cell reversals in 1-D straight channels (Hadjitheodorou et al., 2021), this suggests that the refractory nature of the retracting edge is at least in part mediated by local myosin II/RhoA activity.

Notably, and consistent with our previous observations in straight channels (Hadjitheodorou et al., 2021), the direction of migration reverses considerably before the two cell edges reach equal levels of Cdc42 activity and myosin II localization (cross-point) (Supplementary Figs. 2A-2D). That is, the cell edge that has more Cdc42 activity at any particular moment in time is not necessarily the driving front. Comparing untreated cells to Y27632-treated cells, the latter exhibit faster Cdc42 and myosin dynamics, with the Cdc42 and myosin cross-points happening earlier than in untreated cells but still trailing behind the directional reversal (Supplementary Figs. 2E-2H).

Taken together, our results suggest that, once a cell has made a directional choice, the losing front may enter a refractory period that requires complete retraction before receptor activation can reprogram the edge to switch back to a protrusion program. The refractory nature of the retracting edge can be dampened by reducing myosin II activity.

## DISCUSSION

Overall, our results indicate that cells respond to receptor inputs in a context-dependent way; that is, they exhibit selective “listening” to environmental cues. At the beginning of the competition, both leading edges are committed fronts, and the system is in a balanced state where both directional options are equally viable (Fig. 6). As the cell extends further, both fronts start to lose “confidence” (deplete their Cdc42 activity) and become receptive to new inputs. We can conceptualize this observation by considering an “energy landscape” of decision making. During the competition, the symmetric state (middle valley) becomes less and less energetically favorable as the cell stretches. At this point, optogenetic stimulation becomes more effective for biasing the outcome. After retraction begins, a retracting edge becomes committed to full retraction and typically retracts all the way back to the cell body before it becomes receptive to new receptor inputs (Fig. 6).

**Figure 6:**
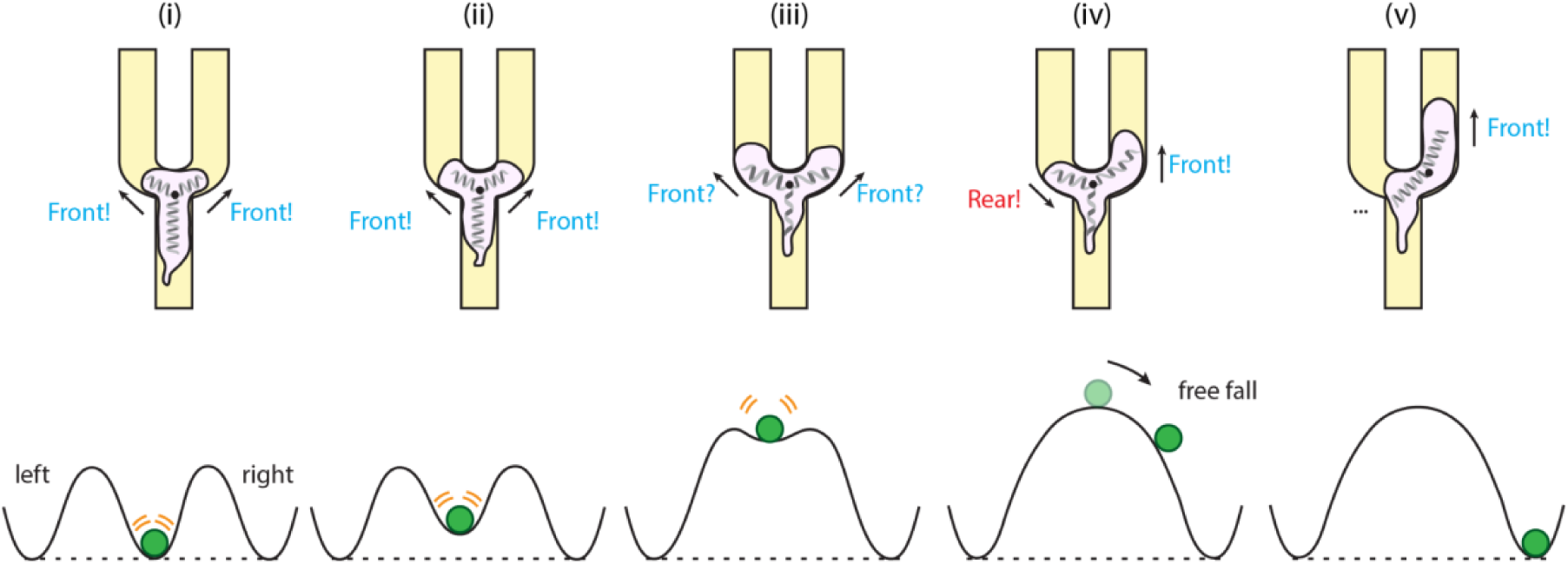
Conceptual model of decision-making in a symmetric directional dilemma. At the onset of the competition, both competing leading edges are committed fronts and the system is in a balanced state (middle valley) where both directional options (left and right) are equally viable (i). Supervised statistical learning reveals no evidence that early pre-existing asymmetries are amplified to help the cells decide which way to turn. Indeed, transient optogenetic receptor activation at this early competition stage is not sufficient to bias the system out of its balanced state. However, as the cell stretches, Cdc42 activity gradually decreases in both competing fronts. Mechanical stretching compromises the self-reinforcing positive feedback loop maintaining Cdc42 activity at the two competing fronts. Conceptually, the symmetric state (middle valley) becomes less and less energetically favorable upon mechanical stretching (ii-iii). Stochastic structural changes at this later stage of competition can be picked up via statistical learning to be statistically correlated with a directional choice. This is also when optogenetic receptor activation to one of the competing fronts is most effective and can bias the cells’ directional choice (iii). After retraction begins (iv), a retracting edge that is committed to full retraction will typically retract all the way back to the apex of the obstacle, comparable to a free fall from an energetic hill, before it becomes receptive to new receptor inputs (v).

An intriguing observation of this work is that both the protrusion speed and the level of Cdc42 activity at a leading front are largely the same with or without optogenetic receptor stimulation (Fig. 4). While the polarity circuit is known to have self-organizing characteristics (Swaney et al., 2010; Wang, 2009), this result was still surprising, given the connections from receptors to front signaling components. However, previous analysis of neutrophil-like cells migrating under agarose found that the front-to-back spatial profiles of Rho GTPases appeared almost indistinguishable irrespective of whether a cell was migrating randomly or in a gradient (Yang et al., 2016). These findings suggest that polarity largely defines the pattern of Cdc42 activity and the other GTPases. Receptor input stimulates and directs the system, but as the system adapts to gradients and inputs, it eventually settles into a more cell-autonomous pattern. Hence, early during the competition the two fronts are fully committed, and the front machinery is maximally engaged. Our optogenetic stimulation does shift the population average behavior, but that is hard to detect at the level of a specific signaling activity like that of Cdc42.

Our data are most consistent with a simple model where a mechanical integrator, cell stretching, contributes to the decline of Cdc42 activity (Fig. 4). The downward trend of the Cdc42 activity at both fronts suggests that cell stretching is impacting the strength of polarity at both fronts, not just at the eventual losing front. A long-standing model in the field, abbreviated as LEGI (local excitation global inhibition), posits that receptor activation generates local protrusive signals as well as a chemical long-range global inhibitor to suppress protrusion at distant sites (Parent and Devreotes, 1999). Under such a model, we would expect to see a decrease in Cdc42 activity at the losing front under optogenetic stimulation, as compared to the activity in the losing front of unstimulated cells undergoing a spontaneous directional choice. However, we were unable to detect any such difference. Instead, the Cdc42 activity measurements shown in Fig. 4 are most consistent with a simple mechanical coupling between the two cell fronts, without the need to invoke an additional LEGI-type chemical signal that communicates the status of one front to its competitor.

From a functional perspective, the context-dependent “listening” to receptor inputs and the refractory nature of the decision-making that our work revealed could represent a particularly efficient way for cells to navigate noisy environments during immune surveillance. Indeed, in cases where the chemical gradient alone might lead cells into an intractable or overly restrictive mechanical path, it is critical that cells have some mechanism to resolve these conflicts without fragmenting or becoming trapped. In these cases, the commitment of a cell front moving in a locally suboptimal direction could provide a solution, allowing the cell to escape and maintain persistent migration. For ultimately reaching their target, this persistence would be more important than short-term accuracy, as the cell can always correct its course later. That is, for an immune cell, it may be better to run now and figure out the details later, rather than to sit in place fixated on the most direct path.

## METHODS

### Cell culture and differentiation

The parental HL60 cell line was a generous gift from Orion Weiner’s lab. In this work, we used the previously published HL60 cell line expressing actin-YFP and MRLC-mApple (Tsai et al., 2019), an HL60 cell line expressing the tdTomato/tdKatushka2 FRET biosensor for Cdc42 and the light-controllable parapinopsin GPCR (Hadjitheodorou et al., 2021), and an HL60 cell line expressing mScarlet-I-tagged Myl9 and parapinospin (Hadjitheodorou et al., 2021). Cells were cultured and differentiated into a neutrophil-like state as previously described (Millius and Weiner, 2009). Briefly, cells were cultured in complete RPMI media (RPMI 1640 plus L-glutamine and 25 mM HEPES media (Gibco RPMI 1640 Medium, HEPES, Thermo Fisher Scientific), supplemented with 10% heat-inactivated fetal bovine serum (hiFBS) (Foundation Fetal Bovine Serum, Gemini Bio, heated in a water bath for 40 min at 56 °C), 100U/mL penicillin, 10 μg/mL streptomycin, and 0.25 μg/mL Amphotericin B (Gibco Antibiotic-Antimycotic, Thermo Fisher Scientific)). Cells were maintained at 37 °C and 5% CO2 in a tissue culture incubator and were passaged every day to maintain a cell density close to 5x10^5^ cells/mL. HL60 differentiation into a neutrophil-like phenotype was achieved by diluting cells in complete RPMI media containing 1.3% DMSO (Acros 61097) at a starting density of 2x10^5^ cells/mL. For all the experiments, only cells differentiated for 6 or 7 days were used.

### Microfluidic device fabrication

Polydimethylsiloxane (PDMS) microfluidic devices were fabricated using standard photolithography and soft lithography approaches. Briefly, we designed two photolithography masks using AutoCAD (Autodesk), one mask for the cell loading channels and dead-end chemoattractant reservoirs, and one mask for the migration channels integrating the bifurcation challenges. Masks were prepared as high-resolution chrome on glass by Front Range Photomasks (Lake Havasu City, Arizona).

To pattern the silicon wafer master-mold, we performed two cycles of spin-coating negative photoresist (SU-8, Microchem, Newton, MA) and exposure to ultraviolet (UV) light though the appropriate masks to produce the migration channels at 3 µm height and cell-loading channels and chemoattractant reservoirs at 200 µm height.

PDMS devices were prepared using the patterned wafer as a replica mold. We mixed PDMS base (Fisher Scientific, Fair Lawn, NJ) with the curing agent at a ratio of 10:1, poured over the wafer, and degassed for at least 1 hour. After baking overnight at 75 °C, PDMS was allowed to cool, cut from the wafer, and inlets and outlets punched using the 1 mm biopsy punch (Harris Uni-Core, Ted Pella, Inc., Redding, CA). Devices were treated with oxygen plasma and bonded to glass-bottom 35 mm γ-irradiated culture dishes with No. 0 coverslips (MatTek Corp., Ashland, MA) by heating to 85 °C for 10 mins on a hotplate.

### Microfluidic device priming

Priming of each device commenced by pipetting 20 μL of an fMLF-based chemoattractant solution through the loading port. For this, fMLF (Sigma Aldrich, stored in stock solutions of 1 μM in DMSO) was diluted in L-15 media (Gibco) containing 20% hiFBS to a final concentration of 100 nM. The device was then placed in a vacuum desiccator connected to house vacuum (27 inHg) for 10 min. Upon removal the device was rested for an additional 10 min, until the chemoattractant filled entirely the straight migration channels, connected orthogonally to the central loading channel. The device was then washed twice, by pipetting 200 μL of 10 μg/mL fibronectin (Sigma Aldrich) diluted in L-15 media containing 20% hiFBS into the loading port. These washing steps removed the chemoattractant from the central loading channel and its passive diffusion from the migration channels into the central loading channel established a chemoattractant gradient. To prevent evaporation, 3 mL of L-15 media were added to the glass-bottomed dish to cover the device.

### Cell staining

To fluorescently label the cell nucleus and microtubules (Figs. 1-2), 3x10^5^ cells were spun down at 200 g for 5 min and resuspended in 1 mL L-15 media (Gibco) containing 1μL SiR-tubulin (Cytoskeleton Inc.) from a stock solution of 1 mM. The stained cell suspension was incubated at 37°C for 45 min. For the last 15 min Hoechst 33324 was added to the cell suspension at a final concentration of 1 μg/mL. Stained cells were spun down at 200 *g* for 5 min and were resuspended in ∼20 μl of L-15. Out of this cell suspension, 10 μL were pipetted into the loading port of the device. The device was then transferred at the microscope stage, for subsequent cell imaging. The microscope was maintained at 37 °C for at least 30 min prior to imaging.

### Retinal preparation and incubation

Preparation of retinal stock solutions was performed in a dark room with red-light sources (Bell et al., 2021). Briefly, 9-cis-Retinal (Sigma Aldrich R5754) was dissolved in argon-purged 200 proof ethanol (Sigma Aldrich) to reach a concentration of 10 mg/mL. Aliquots were stored at -80 °C in small amber glass tubes (Sigma Aldrich). 0.1 g of Bovine Serum Albumin (BSA), Fraction V—Low-Endotoxin Grade (Gemini Bio 700-102P) was mixed in 10 mL of L-15 media (Gibco), to prepare a 1% BSA solution. In darkness, 10 μL of retinal stock was diluted to a working concentration of 10 μg/mL by gradually adding the 1% BSA solution until all 10 mL were used. The final retinal solution was mixed overnight at 4 °C on a rocker in total darkness. Final retinal solutions were used for experiments within 2 days after preparation.

For the stimulation assays, 3 x 10^5^ differentiated HL60 cells were spun down at 200 g for 5 min and re-suspended in 1 mL of retinal solution to incubate at 37 °C for 1 h. This incubation and all remaining cell handling happened in darkness with red-light sources. Incubated cells were spun down at 200 g for 10 min and were resuspended in ∼ 20 μL of L-15. Out of this cell suspension, 10 μL were pipetted into the loading port of the device. The device was then transferred on the microscope at 37 °C, where the cells would be imaged.

### Pharmacological treatments

To explore the role of microtubules in decision-making (Supplementary Fig. 1) as well as the role of myosin II/RhoA in creating the refractory nature of the retracting losing front to receptor inputs (Fig. 5 and Supplementary Fig. 2) we pre-treated cells with either 50 μM nocodazole (Sigma Aldrich) or 10 μM Υ-27632 (Sigma Aldrich) for the last 30 min of the staining or retinal incubation, respectively. Cells were spun down at 200 g for 5 min and were re-suspended in ∼ 20 μL of L-15 containing the same final drug concentration. Of this cell suspension, 10 μL were loaded in a device that was pre-treated with the same drug concentration. The pharmacological compound was included in both the chemoattractant solution as well as the washing solution used during device priming, in the same final concentration. For all pharmacological treatments DMSO was used as a vehicle. We performed control experiments with 0.2% DMSO to match the maximum final DMSO concentration in the pharmacological treatments. We found no significant differences between untreated and DMSO-treated cells.

### Fluorescence microscopy for phase contrast and intracellular structures

Multi-channel time-lapse sequences of epifluorescence (to image the fluorescently-tagged cytoskeletal structures) and phase contrast images (to image the cell shape) were acquired using an inverted Nikon Eclipse Ti2 with an EMCCD camera, Andor iXon3 (Andor Technologies). A Lambda XL (Sutter Instruments) light source was used for epifluorescence illumination. A filter wheel in the light source permits rapid selection of excitation wavelength. Images were captured every 3 s using either a 20x 0.75 NA Plan Apo air objective (for Fig. 1B) or a 100x 1.45 NA oil immersion objective (for Figs. 1C-1D, Fig. 2 and Supplementary Fig. 1). An acrylic microscope incubation chamber (Haisen Tech) was used to maintain the microscope at 37 °C during the entire imaging session. Cells were imaged using sequential phase contrast and epifluorescence illumination. To capture single cell actin-myosin dynamics we used a dual filter cube (Chroma, #59022) to capture eGFP and mCherry emission by sequential excitation with 470/40 nm and 560/40 nm filters respectively. To record Hoechst-stained cell nucleus and SiR-tubulin-stained cell microtubules signals we used standard DAPI and Cy5 (Chroma, #49006) filter cubes, respectively. Micro-Manager was used to operate all microscopy equipment (Edelstein et al., 2014).

### Microscope configuration for Cdc42 TomKat FRET sensor and Myl9 imaging

Fluorescence microscopy for the Cdc42 TomKat FRET sensor and the Myl9 imaging was performed on a Nikon Eclipse Ti inverted microscope with a XLED1 LED (Excelitas) light source for epifluorescence illumination. Images were acquired every 3 s on two Andor Zyla 4.2 sCMOS cameras, using a Cairn TwinCam LS image splitter equipped with a 605 nm edge wavelength dichroic mirror (Chroma ZT594rdc) and emission filters to allow simultaneous imaging. The Cdc42 TomKat FRET imaging was performed as previously described (Bell et al., 2021). For myosin Myl9 imaging experiments, we used a 405/488/561/640 nm quad band filter cube (Chroma 91032) that allowed rapid, sequential 407 nm optogenetic stimulation, and simultaneous imaging of mScarlet-I-Myl9 (∼ 561 nm excitation), and cytoplasmic iRFP (∼ 640 nm excitation). Bandpass emission filters were used to eliminate bleedthrough of iRFP into the myosin channel. Some bleedthrough of mScarlet signal into the iRFP channel was unpreventable and tolerated to allow simultaneous imaging. Rapid and precise stage movements were achieved using an ASI stage (MS- 2000 Flat-top) equipped with linear encoders. The microscope was controlled by custom-built MATLAB R2015a software (MathWorks) interfaced with Micro-Manager (Version 1.4.23), allowing automated and highly reproducible experimental conditions for cell stimulation and time-lapse microscopy protocols. Cells were imaged with a 60x oil immersion objective (Nikon Apochromat, 1.49 NA) and were maintained at 37 °C using a temperature and humidity control unit (OkoLab Microscope Lexan Enclosure). Each experiment was terminated within 2 h of imaging.

### Stimulation assays

Cell stimulation and imaging was performed using a custom-build MATLAB R2015a interface for Micro-Manager (Version 1.4.23). Opsin stimulation was performed using a 407 nm laser (Coherent Cube) with a custom fiber coupling inserted in a FRAP port on the microscope. Both the TomKat dichroic and the dichroic used for myosin imaging can pass 407 nm light, enabling rapid FRAP stimulation. To focus the FRAP module, HL60 cells expressing either the TomKat FRET sensor or the mScarlet-I-Myl9 were loaded in a microfluidic device and imaged, to determine the appropriate focal plane. Cells were thereafter selected, zapped (setting the laser to 40 ms exposure and 10 % power) and imaged using the FRAP channel. The *x*-*y* translation knobs of the FRAP module were adjusted to bring the FRAP spot near the center of the image (around 512 pixel x 512 pixel on a 1024 pixel x 1024 pixel image, where 1 pixel is 0.21 µm). Thereafter, the *z*- adjustment knob was used to focus the FRAP spot into a tight approximately gaussian-shaped spot. Cell-stimulation experiments were conducted using ∼1.8 µW power, as measured at the objective using a Thorlabs handheld optical power meter (PM100D) and a microscope slide power sensor (S170C). For each experimental session, we collected pictures of the FRAP spot and averaged them to identify the pixel with the maximum FRAP spot intensity. The *x*-*y* coordinates of the maximum intensity pixel were saved and later used to dynamically translate the stage, to align the maximum activation spot with the desired sub-cellular stimulation location.

Each lane of the microfluidic device was organized into a grid of *x*-*y* coordinates that were imaged consecutively. Cells were segmented in real time using a minimum fluorescence threshold and minimum and maximum size thresholds. The target point was computed by determining the left front (in the competition assays) or the retracting/losing front (in the reversal assays). Determination of the cell front relied on extracting the cell body boundary from the binary mask and determining the point in the cell perimeter as specified by a target angle (for us 0^o^ in respect to the original orientation of cell movement). For continuous and late stimulation experiments, UV stimulation and imaging were alternated until the imaging period was over. For the early stimulation assays we administered a total of 5 pulses of light. For all assays we maintained the same time interval (3 s) for imaging and stimulation. Alternating between imaging and stimulation, results in turning the receptor off with each image acquisition and then rapidly back on again with the following stimulation pulse, as parapinopsin is a Giα-family coupled GPCR that is activated by UV light and inactivated by orange light (> 530 nm) leading to rapid activation and deactivation cycles. Thus, our stimulation assays are pulsed.

### Camera and illumination corrections for Cdc42 TomKat FRET and Myl9 imaging

Raw images were corrected for the camera dark-state noise, for differences in the camera chip sensitivity, and for dust in the light path as previously described (Bell et al., 2021). In addition, a gradient in apparent FRET ratio activity was empirically observed from the top to bottom of the TomKat FRET sensor images, due to imperfections in the light path. We corrected for this gradient by developing a “ratio correction image”. For that, images of unstimulated Cdc42 TomKat FRET sensor HL60 cells loaded in microfluidic channels were collected systematically with different stage positions and the same 60x objective so that at least one cell was imaged on every portion of the camera sensor. FRET ratios were computed using our standard analysis pipeline for each image and images were assembled into a 3D image stack. To generate a single representative full-field FRET image, we computed the median of the FRET ratio over the stack (including only pixels corresponding to cells). To reduce local variability, we smoothed this image by taking the median over each 24 pixel x 24 pixel block and smoothed using a gaussian filter (sigma=5). Finally, we resized the result to generate a ratio correction image with the same dimensions as the input image. We applied the correction by dividing the FRET ratio images by the computed ratio correction image. This gradient correction was not necessary for the Myl9 imaging.

### Cell segmentation and image background subtraction for Cdc42 TomKat FRET and Myl9 imaging

Raw Cdc42 TomKat FRET pair images were registered using the coordinate-mapping strategy described (Bell et al., 2021). Cell segmentation and image background subtraction for the Cdc42 TomKat FRET images and the Myl9 images were performed as described before (Hadjitheodorou et al., 2021). Briefly, cell segmentation was performed on the sum of the aligned FRET donor and acceptor images, to improve signal to noise ratio. We first conservatively defined background and cell object pixels and then determined a background image using the median intensity of background pixels in the local neighborhood of each pixel. We then subtracted the background image from the sum image. Object edges were enhanced by first smoothing the image using a broader gaussian filter (sigma=5), and then subtracting the smoothed image from the original image. Finally, the cell object binary masks were determined through the Otsu’s threshold method. For each FRET donor and acceptor image we subtracted the background as defined through the above segmentation strategy. Pixels not included in the cell mask were defined as not a number (NAN), to eliminate them from downstream analysis. The FRET ratio image was calculated as FRET acceptor divided by FRET donor. We used a similar strategy to register the mScarlet-I-Myl9 and cytoplasmic iRFP pair images for the myosin cell line and to subtract the background.

### Statistical learning

#### Pre-processing

Precise segmentation of cellular boundaries was critical in building the statistical learning models. Empirically, we found that manual user correction was necessary to ensure the quality of segmentation of the phase contrast images. We developed a semi-automatic workflow using the magnetic lasso and quick selection tool in Adobe Photoshop CS6 (Adobe Systems Inc.) to enable fast manual segmentation.

After defining background and cell object pixels, we calculated the median intensity of background pixels in the local neighborhood of the cell for all fluorescence images. The background was subtracted from the raw fluorescence images and fluorescence intensity was normalized across movies. All images were centered at the cell centroid and cropped to a consistent size (601 pixels x 601 pixels).

#### Feature extraction

For each cell decision-making movie, we found the image frames from when the cell first contacted the obstacle until retraction initiation. Retraction initiation was defined as the first frame where we identified a retraction area of the losing front (*i.e.*, area difference > 0). For these frames we used the open software CellProfiler 3.0 to extract an array of features from the binary masks and the fluorescence images using a custom-made image analysis pipeline. Specifically, we extracted features based on morphology (size and shape from the *MeasureObjectSizeShape* module) and intensity-based parameters (intensity statistics, texture, and granularity within segmented cells, from the *MeasureObjectIntensity*, *MeasureObjectRadialDistribution*, *MeasureTexture* and MeasureImageGranularity modules). A full list of the morphological parameters is provided in Supplementary Table 1. Our pipeline returned a total of ∼700 features for each time point for further processing.

#### Post-processing

Different cells make their decisions at different rates (Supplementary Fig. 1A). To facilitate comparisons across the large population of individual cells, we filtered the data to include only events whose time from contact with the obstacle until retraction initiation was within one standard deviation of the population average (45 s ± 15 s). To synchronize the filtered decision-making events, we worked on a rescaled time, interpolating the feature values extracted from the frames of each cell decision-making movie for 15 time points between contact with the obstacle and retraction initiation, so that on average the time between two consecutive interpolated time points is ∼ 3 s (equal to the frame rate of the microscopy data). Feature interpolation was carried out via fitting a cubic smooth spline with the smoothing parameter set at the default value of 0.5 (where 0 fits a straight line and 1 gives a total fit) for each cell and each feature over time.

#### Feature selection

Considering the small number of training samples in relation to the large number of extracted features (696 features extracted through CellProfiler), we predicted the binary outcome of directional decision-making using the lasso (L1 penalty) logistic regression. The lasso enables fitting a logistic regression even when the number of features exceeds the number of observations, regularizes the parameters of the logistic regression to reduce model variance, and discards features to maintain only the most predictive ones. For each dataset examined, we trained on 67 cells and tested the classifier on an independent test set of 33 cells with known labels (1 for right turning cells and 0 for left). Training and testing were performed for each time point separately. For each time point, all features were standardized by subtracting the cell average and dividing by the standard error of the mean. We cross-validated the model by implementing the cv.glmnet function with nfold=10 from the glmnet package (Friedman et al., 2010) in R. The tuning parameter λ was chosen by the 10–fold cross validation as the λ for the best model path. We used the selected model to evaluate the misclassification error in the test set. The CellProfiler features and the lasso feature coefficients across interpolated time points are presented in Supplementary Table 1. In addition to using the lasso on features extracted through CellProfiler, we also implemented the lasso on cumulative feature means as well as feature slopes computed over consequent interpolated time points, however these strategies did not improve the prediction.

We also implemented an elastic net regression and a sparse group lasso regression (Supplemental Figures 1C-1D). Elastic Net was implemented using the glmnet package; Supplementary Figure 1 illustrates the predictive accuracy, which is equal to (1-misclassification error) %, of the statistical models across interpolated time points with α=0.8 (where α=0 is ridge L2 penalty and α=1 is lasso L1 penalty). For the sparse group lasso strategy, we formed 696 feature groups, one for each feature over the 15 interpolated time points. We then applied sparse group lasso and were left with 20 groups with non-zero features. Finally, using just these 20 features we used an elastic net model with α=0.8 at each time independently and calculated the misclassification error in the test set. Even though this strategy allows the earlier time points (closer to the cell contacting the obstacle) to be informed by later ones (closer to directional decision), it showed comparable results with that of the initial lasso (Supplemental Figures 1C-1D).

### Image analysis

#### Quantification of Cdc42 activity and myosin localization at each edge

Cell front areas were defined as the 400 pixels closest to the cell front edge. The 400 pixels enabled us to capture the entire penetration depth of the Cdc42 activity at the competing cell fronts in this channel geometry. The cell front edge point was defined as the cross section point between the midline of the channel and the cell mask in each arm of the bifurcation. We computed the Cdc42 sensor activity as the sum of the acceptor signal divided by the sum of the donor signal in that area (Figs. 4B, 4E, 4G, 4H, 5F and Supplementary Figs. 2B, 2F). Cell front Cdc42 activities were corrected for imaging-induced photobleaching using an empirically determined linear, multiplicative photobleaching correction. The photobleaching correction was computed by fitting a linear rate for the decrease in measured FRET ratio for the same Cdc42 sensor in an equal sized cell front region in well polarized cells migrating persistently without experimental perturbation in straight microfluidic channels (Hadjitheodorou et al., 2021).

We used the same strategy to identify the front of Myl9-expressing cells, so that we can directly compare the dynamic re-localization of Cdc42 activity and myosin II (Supplementary Fig. 2). To analyze the relative subcellular distribution of myosin, we computed the normalized myosin at the front as the sum of the Myl9 signal in each area of interest divided by the sum of the Myl9 signal in the entire segmented cell body (Supplementary Figs. 2D, 2H). No photobleaching correction was necessary for the Myl9 signal, as photobleaching was negligible compared to the dynamic range of the Myl9 signal.

#### Cell speed, length and area quantification

To calculate the protrusion and retraction speed, we compared the 400 pixel masks every other frame, and marked the newly protruded regions and the newly retracted regions. The regions were summed up and divided by the time step in between frames to obtain the instantaneous protrusion speed and retraction speed, hence the units of speed are μm^2^/min (Figs. 4A, 4D, 5E and Supplementary Figs. 2A, 2C, 2E, 2G). Cell length was approximated via computing the Y-shaped skeleton of the binary cell mask and summing up its length segments, while cell area was calculated by summing the total number of pixels of the computed binary cell mask. Relative changes in length and area were plotted by dividing the cell length and area for each cell by its respective length and area at either the time of contact with the obstacle (Figs. 4C, 4F) or at the time when the two competing fronts had clearly separated from one another (*i.e.,* when both had extended at least 2.2 µm past the apex) (Figs. 4G, 4H).

#### Temporal cross correlation analysis

Relative change in cell length, cell stretching, was computed as described directly above, relative to cell length at the time when the two competing fronts had clearly separated from one another. The difference in front Cdc42 activity between the competing fronts was determined by subtracting the Cdc42 activity of the losing front from the Cdc42 activity of the winning front and taking the absolute value of that difference. To reduce noise, we examined traces starting at the time when the two competing fronts had clearly separated from one another and ending at the time of big retraction. In addition, we kept cells with meaningfully long traces (10 time frames, corresponding to 30 s), and excluded cells with shorter traces. These parameter traces were smoothed via a local regression using weighted linear least squares and a 2^nd^ degree polynomial model (“loess” or locally weighted smoothing in MATLAB). We used a span of 10% of the total number of data points for this smoothing. Temporal cross-correlations were then computed as a function of offset in time between cell stretching and absolute difference in Cdc42 activity between the winning and the losing cell front using Pearson’s correlation.

#### Stratification of fast and slow responding reversing cells

Computing the edge point of the retracting front as described above, we calculated the euclidean distance of that point relative to the obstacle apex point of the obstacle at two times: i) when the stimulation protocol was initiated (*i.e.* when a big retraction was detected) and we plot that on the *x*-axis of Fig. 5D, and ii) when the retracting edge had reached its minimum distance from the obstacle apex before reprotrusion commences (reprotrusion is defined as protrusion area more than 7.2 µm^2^) and we plot that on the *y*-axis of Fig. 5D. For each point, we then took the *y* coordinate and divided it by the *x* coordinate to get one number for each cell. That number should be near zero for a cell that retracts fully and near 1 for a cell that immediately reverses. We called cells for which *y*/*x* < 0.5 slow responders, and the rest fast responders.

### Statistical analysis

Day-to-day and experiment-to-experiment variation for categorical data was consistent with Poisson counting statistics and the observed standard deviations were reasonably explained by the binomial model in each case. For quantitative signaling, speed, and localization data, cell-to-cell variability was the dominant source of variability. Given the limited and variable number of cells that could be analyzed in each experiment, we believe that individual cells represented the most relevant independent unit for statistical analysis. In addition, we performed control experiments on each day to avoid potential bias. Based on these considerations, we pooled data from all independent experiments performed. We indicate the collective number of cells per condition in the Figure Legends. All measurements were taken using independent cells. Statistical parameters and significance are reported in the Figures and the Figure Legends. Data are determined to be statistically significant when *p* < 0.05 by either two-sided Wilcoxon rank sum test or two-sided Fisher exact test (“ranksum” or “fishertest” in MATLAB, respectively).

To depict the percentage of cells across the different assays (Fig. 1B, 3C, and 5B), we represented data with bar plots showing the cumulative mean value and an error bar which represents the standard deviation assuming a binomial distribution around the cumulative mean, pooling data from all independent experiments.

For the statistical analysis of the time from front contact to (big) retraction (Fig. 4I and Supplementary Fig. 1F), the cell speed (Fig. 5E) and the subcellular localization of Cdc42 activity (Fig. 5F), we used violin plots pooling all cells that passed our pre-processing standards of masking and tracking, from all independent experiments. In each violin plot the dashed line represents the median of the distribution and the dotted lines the 25% and 75% quartiles of the distribution. To compare across conditions or cell groups, we performed non-parametric two-sided Wilcoxon rank sum tests.

### Movie processing

Supplementary Movies 1-3 show background subtracted signals, as described above. For the Supplementary Movies 4-9, each donor and acceptor image was smoothed using a gaussian filter (sigma=2), after subtracting the background as previously described. This smoothing step was only applied to movies and not for any other analysis. The FRET ratio image was calculated as FRET acceptor divided by FRET donor (using the smoothed images). The Cdc42 activity ratio for each cell was a bit different and since the relative activity across each cell is most informative, we chose to show cells in Supplementary Movies 4-9 with a FRET range from the 1^st^ percentile to the 99^th^ percentile of the FRET ratio images of each movie.

### Plotting

Plotting was performed using MATLAB R2018a (MATHWORKS) and GraphPad Prism 9.

## ACKNOWLEDGMENTS

We thank Tony Y.-C. Tsai, Effie Bastounis and Nathan Belliveau for insightful discussions and comments on the manuscript. This work was supported by the Howard Hughes Medical Institute (J.A.T.), and an NIH Director’s New Innovator Award (DP2 HD094656) to S.R.C. A.H. was a Stanford Bio-X Bowes Fellow, an Onassis Foundation Scholar, and was supported by the Foundation for Education and European Culture (IPEP), the A.G. Leventis Foundation and the Attica Tradition Foundation. Microfabrication work was supported by NIG grant GM092804 to D.I. G.R.R.B. thanks the National Science Foundation Graduate Research Fellowship (Grant Number 1650042) for support.

This article is subject to HHMI’s Open Access to Publications policy. HHMI lab heads have previously granted a nonexclusive CC BY 4.0 license to the public and a sublicensable license to HHMI in their research articles. Pursuant to those licenses, the author-accepted manuscript of this article can be made freely available under a CC BY 4.0 license immediately upon publication.

## AUTHOR CONTRIBUTIONS

A.H., G.R.R.B., S.R.C. and J.A.T. designed the experiments, A.H. performed the experiments and analyzed the data, R.T. and A.H. developed the statistical learning models, A.H., G.R.R.B. and S.R.C. created software for data analysis and automating experiments, G.R.R.B. and S.R.C. developed the optogenetic and imaging tools, F.E. and D.I. designed the microfluidic devices, A.H., S.R.C. and J.A.T. interpreted the data, A.H., S.R.C. and J.A.T. wrote the manuscript. All authors read and approved the manuscript. R.T. was supported by the National Institutes of Health (5R01 EB001988-16) and the National Science Foundation (19 DMS1208164).

## DATA AVAILABILITY STATEMENT

The data that support the findings of this study are available from the corresponding authors upon request.

## CODE AVAILABILITY STATEMENT

The code we developed for automating experiments and data analysis is available from the corresponding authors upon request.

---

**Supplementary Figure 1:**
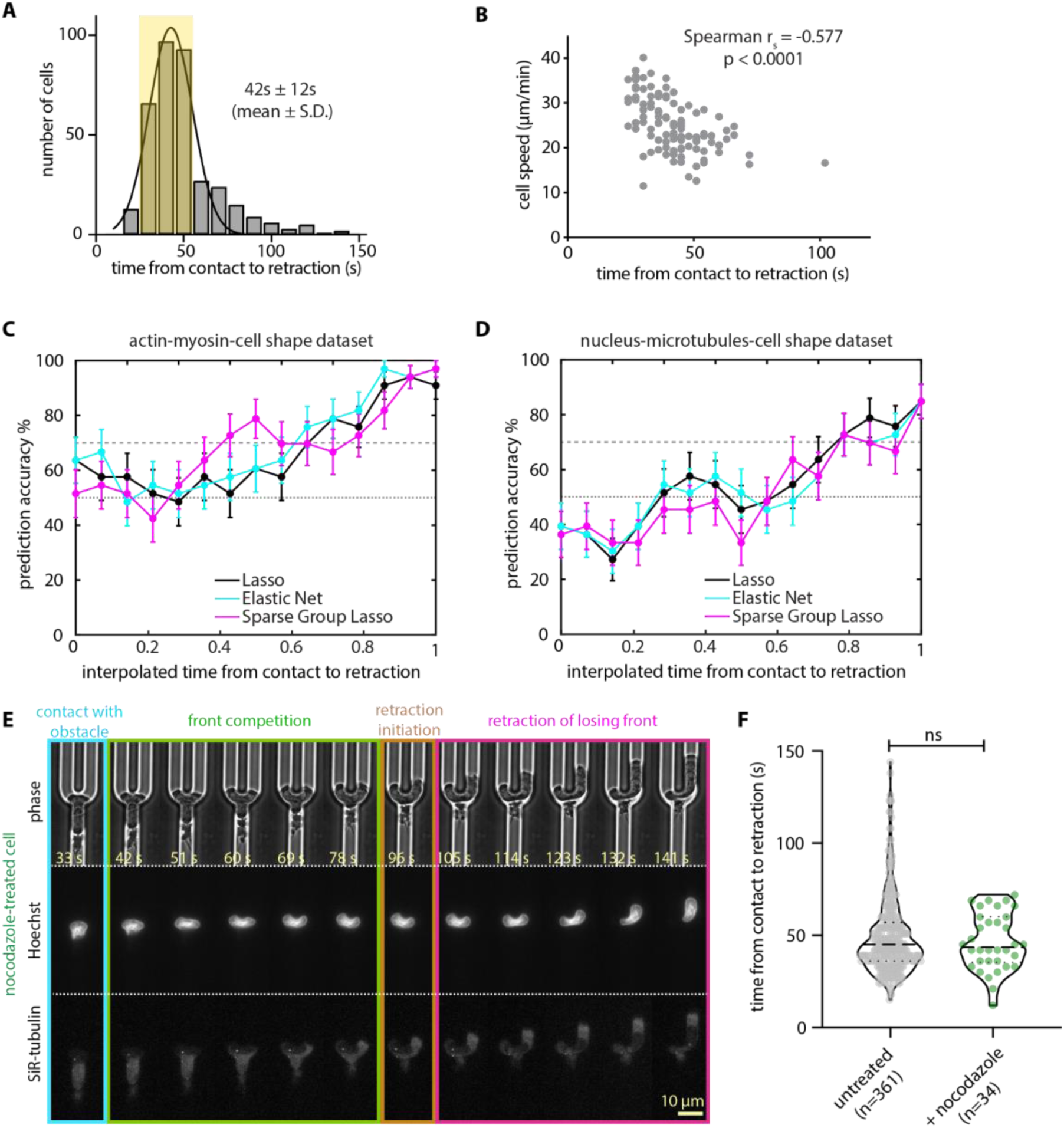
Timing and predictability of cellular decisions in bifurcation devices. **(A)** Histogram of time from contact with the obstacle until retraction initiation for 361 cells migrating inside symmetrically bifurcating microfluidic channels. Gaussian fit yields a mean of 42 s and a S.D. of 12 s. Cells whose decision time was within one standard deviation of the mean were retained for further analysis (yellow box). **(B)** Scatter plot of average cell centroid speed before contact with the obstacle vs. time from contact with the obstacle until retraction initiation for 109 cells. Spearman correlation yields rs = -0.577, p < 0.0001. **(C-D)** Test set prediction accuracy of statistical models leveraging information contained in binary masks and fluorescence images of actin and myosin **(C)** and in binary images and fluorescence images of nucleus and microtubules **(D)** across interpolated time points between contact with the obstacle and retraction initiation. **(E)** Representative live-cell imaging snapshots of single HL60 cells treated with nocodazole to depolymerize the microtubules (red channel) and stained with Hoechst to tag the nucleus (cyan) during symmetric decision-making (Supplementary Movie 3). Images were captured every 3 s and subsampled for illustration purposes. Scale bar: 10 μm. **(F)** Violin plot of time from contact with the obstacle to retraction initiation for 361 untreated and 34 nocodazole-treated cells; *p*-value of two-sided Wilcoxon’s rank-sum test (ns: *p* > 0.05).

**Supplementary Figure 2:**
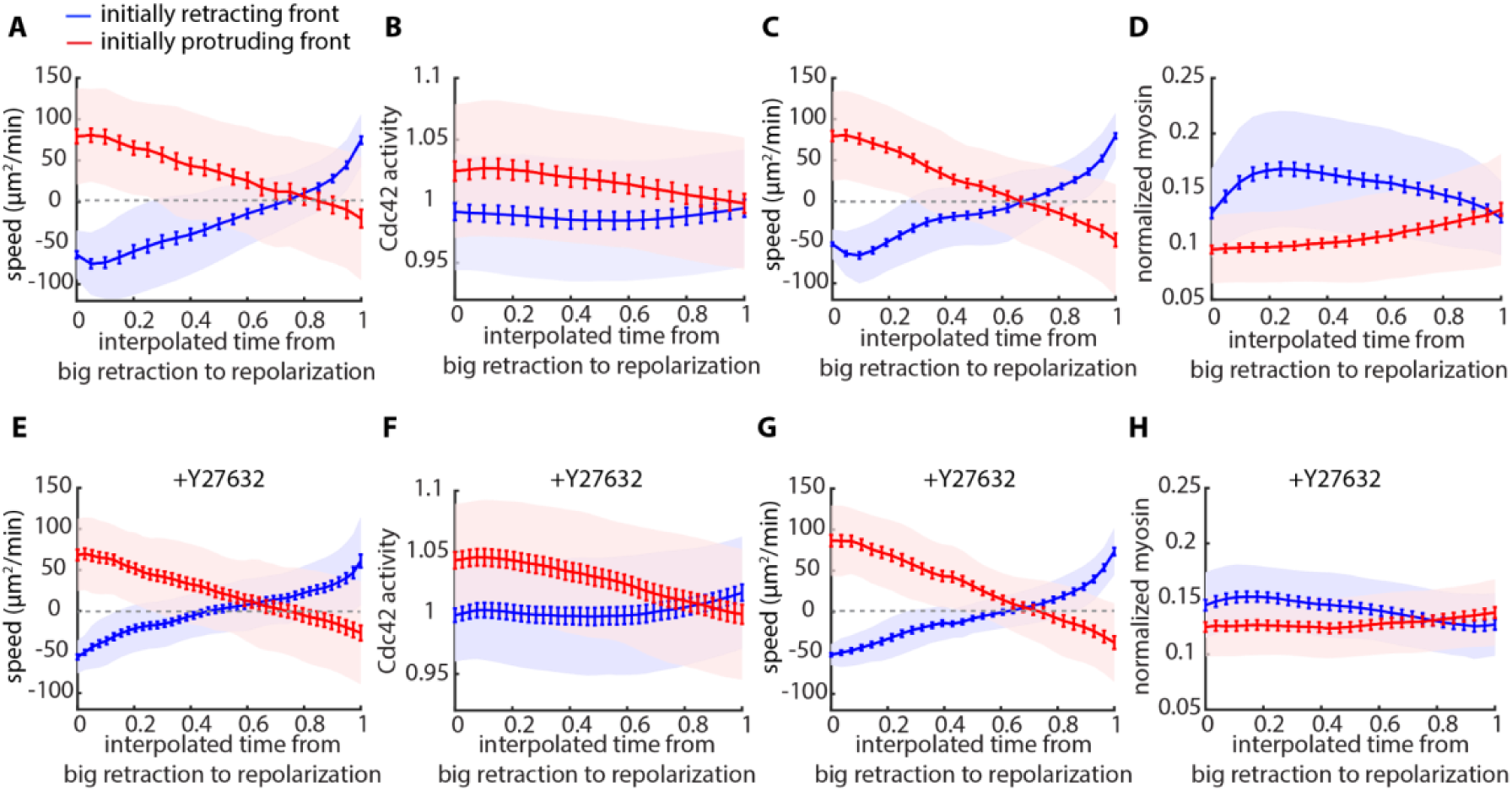
Cdc42 activity and myosin II polarity reverse after directional change. Mean speed **(A)** and magnitude of Cdc42 activity **(B)** of the initially retracting (stimulated, blue) and initially protruding (red) edges for 47 cells expressing parapinopsin and Cdc42 FRET sensor that successfully reversed under the continuous stimulation reversal assay (lines: means, shaded regions: SD, error bars: mean value ± SE). Mean speed **(C)** and normalized myosin intensity **(D)** of the initially retracting (stimulated, blue) and protruding (red) edges for 70 cells expressing parapinopsin and Myl9 that successfully reversed under the continuous stimulation assay (lines: means, shaded regions: SD, error bars: mean value ± SE). Mean speed **(E)** and magnitude of Cdc42 activity **(F)** of the initially retracting (stimulated, blue) and initially protruding (red) edges for 27 Y27632-treated cells expressing parapinopsin and Cdc42 FRET sensor that successfully reversed under the continuous stimulation assay (lines: means, shaded regions: SD, error bars: mean value ± SE). Mean speed **(G)** and normalized myosin intensity **(H)** of the initially retracting (stimulated, blue) and initially protruding (red) edges for 37 Y27632-treated cells expressing parapinopsin and Myl9 that successfully reversed under the continuous stimulation assay (lines: means, shaded regions: SD, error bars: mean value ± SE).

## Additional Files

**Supplementary Table 1**: Tables of the analyzed CellProfiler features extracted from HL-60 cells undergoing symmetric decision-making. This Excel file contains two sheets. The first sheet contains the Lasso coefficients for the features extracted from the dataset composed of phase contrast images, along with fluorescence images of actin-YFP and MRLC-mApple. The second sheet contains the contains the Lasso coefficients for the features extracted from the dataset composed of phase contrast images, along with fluorescence images of the cell nucleus (Hoechst) and microtubules (SiR-tubulin).

**Supplementary Movie 1: Decision-making of an HL60 cell facing a symmetric directional dilemma.**

Movie of an HL60 cell expressing actin-YFP and MRLC-mApple. On the left panel we show the overlay of the phase contrast image, actin and myosin fluorescence and next to that the individual microscopy channels (in grayscale). Images were captured every 3 s. Scale bar: 10 μm.

**Supplementary Movie 2: Decision-making of an HL60 cell facing a symmetric directional dilemma.**

Movie of an HL60 cell stained with Hoechst to tag the cell nucleus and with SiR-tubulin to tag the microtubules. On the left panel we show the overlay of the phase contrast image, nuclear and microtubules’ fluorescence and next to that the individual microscopy channels (in grayscale). Images were captured every 3 s. Scale bar: 10 μm.

**Supplementary Movie 3: Decision-making of a nocodazole-treated HL60 cell facing a symmetric directional dilemma.**

Movie of a nocodazole-treated HL60 cell stained with Hoechst to tag the cell nucleus and with SiR- tubulin to tag the microtubules. On the left panel we show the overlay of the phase contrast image, nuclear and microtubules’ fluorescence and next to that the individual microscopy channels (in grayscale). Images were captured every 3 s. Scale bar: 10 μm.

**Supplementary Movie 4: A cell turns left under continuous optogenetic stimulation at the left competing front.**

Movie of a continuous stimulation competition assay for an HL60 cell expressing parapinopsin and the Cdc42 FRET sensor. Initially, the cell migrated without experimental perturbation and after extending two competing fronts clearly separated from one another (*i.e.,* when both had extended at least 2.2 µm past the apex), we administered continuous optogenetic stimulation at its left competing front (magenta circle). On the left we show the registered sensors (in grayscale) and on the right the computed Cdc42 sensor activity (FRET ratio). Images were captured every 3 s. Scale bar: 25 μm.

**Supplementary Movie 5: A cell not responding to early optogenetic stimulation at the left competing front.**

Movie of an early stimulation competition assay for an HL60 cell expressing parapinopsin and the Cdc42 FRET sensor. Initially, the cell migrated without experimental perturbation and after extending two competing fronts clearly separated from one another (*i.e.,* when both had extended at least 2.2 µm past the apex), we administered 5-pulses of optogenetic stimulation at its left competing front (magenta circle). On the left we show the registered sensors (in grayscale) and on the right the computed Cdc42 sensor activity. Images were captured every 3 s. Scale bar: 25 μm.

**Supplementary Movie 6: A cell turns left under late optogenetic stimulation at the left competing front.**

Movie of a late stimulation competition assay of an HL60 cell expressing parapinopsin and the Cdc42 FRET sensor. Initially, the cell migrated without experimental perturbation and after extending two competing fronts have extended at least 6.6 µm past the apex we administered continuous optogenetic stimulation at its left competing front (magenta circle). On the left we show the registered sensors (in grayscale) and on the right the computed Cdc42 sensor activity. Images were captured every 3 s. Scale bar: 25 μm.

**Supplementary Movie 7: A cell reversing its direction of motion under continuous optogenetic stimulation at the retracting/losing front.**

Movie of a successful continuous stimulation reversal of an HL60 cell expressing parapinopsin and the Cdc42 FRET sensor. The cell migrates and decides a direction of turning without experimental perturbation. Upon identifying a big retraction, we administer continuous stimulation (magenta circles) at the retracting/losing front. On the left we show the registered sensors (in grayscale) and on the right the computed Cdc42 sensor activity. Images were captured every 3 s. Scale bar: 25 μm.

**Supplementary Movie 8: A cell not reversing its direction of motion under early optogenetic stimulation at the retracting/losing front.**

An unsuccessful early stimulation reversal of an HL60 cell expressing parapinopsin and the Cdc42 FRET sensor. The cell migrates and decides a direction of turning without experimental perturbation. Upon identifying a big retraction, we administer 5-pulses of optogenetic stimulation (magenta circles) at the retracting/losing front. On the left we show the registered sensors (in grayscale) and on the right the computed Cdc42 sensor activity. Images were captured every 3 s. Scale bar: 25 μm.

**Supplementary Movie 9: A cell reversing its direction of motion under late optogenetic stimulation at the retracting/losing front.**

A successful late stimulation reversal of an HL60 cell expressing parapinopsin and the Cdc42 FRET sensor. The cell migrates and decides a direction of turning without experimental perturbation. We let the cell retract back to the apex of the obstacle and then began repetitively stimulating the previously retracting/losing front (magenta circles). On the left we show the registered sensors (in grayscale) and on the right the computed Cdc42 sensor activity. Images were captured every 3 s. Scale bar: 25 μm.

## REFERENCES

Ambravaneswaran, V., Wong, I.Y., Aranyosi, A.J., Toner, M., and Irimia, D. (2010). Directional decisions during neutrophil chemotaxis inside bifurcating channels. Integr Biol (Camb) 2, 639– 647.

Bell, G.R.R., Rincón, E., Akdoğan, E., and Collins, S.R. (2021). Optogenetic control of receptors reveals distinct roles for actin- and Cdc42-dependent negative signals in chemotactic signal processing. Nature Communications 12, 6148.

Belotti, Y., McGloin, D., and Weijer, C.J. (2020). Analysis of barotactic and chemotactic guidance cues on directional decision-making of Dictyostelium discoideum cells in confined environments. PNAS 117, 25553–25559.

Carpenter, A.E., Jones, T.R., Lamprecht, M.R., Clarke, C., Kang, I.H., Friman, O., Guertin, D.A., Chang, J.H., Lindquist, R.A., Moffat, J., et al. (2006). CellProfiler: image analysis software for identifying and quantifying cell phenotypes. Genome Biology 7, R100.

Edelstein, A.D., Tsuchida, M.A., Amodaj, N., Pinkard, H., Vale, R.D., and Stuurman, N. (2014). Advanced methods of microscope control using μManager software. Journal of Biological Methods 1, e10–e10.

Ellett, F., Marand, A.L., and Irimia, D. (2022). Multifactorial assessment of neutrophil chemotaxis efficiency from a drop of blood. J Leukoc Biol.

Etienne-Manneville, S., and Hall, A. (2002). Rho GTPases in cell biology. Nature 420, 629–635.

Friedman, J., Hastie, T., and Tibshirani, R. (2010). Regularization Paths for Generalized Linear Models via Coordinate Descent. J Stat Softw 33, 1–22.

Hadjitheodorou, A., Bell, G.R.R., Ellett, F., Shastry, S., Irimia, D., Collins, S.R., and Theriot, J.A. (2021). Directional reorientation of migrating neutrophils is limited by suppression of receptor input signaling at the cell rear through myosin II activity. Nat Commun 12, 6619.

Hind, L.E., Vincent, W.J.B., and Huttenlocher, A. (2016). Leading from the back: the role of the uropod in neutrophil polarization and migration. Dev Cell 38, 161–169.

Houk, A.R., Jilkine, A., Mejean, C.O., Boltyanskiy, R., Dufresne, E.R., Angenent, S.B., Altschuler, S.J., Wu, L.F., and Weiner, O.D. (2012). Membrane Tension Maintains Cell Polarity by Confining Signals to the Leading Edge during Neutrophil Migration. Cell 148, 175–188.

Jilkine, A., Marée, A.F.M., and Edelstein-Keshet, L. (2007). Mathematical model for spatial segregation of the Rho-family GTPases based on inhibitory crosstalk. Bull Math Biol 69, 1943– 1978.

Kopf, A., Renkawitz, J., Hauschild, R., Girkontaite, I., Tedford, K., Merrin, J., Thorn-Seshold, O., Trauner, D., Häcker, H., Fischer, K.-D., et al. (2020). Microtubules control cellular shape and coherence in amoeboid migrating cells. J Cell Biol 219.

Lämmermann, T., Renkawitz, J., Wu, X., Hirsch, K., Brakebusch, C., and Sixt, M. (2009). Cdc42-dependent leading edge coordination is essential for interstitial dendritic cell migration. Blood 113, 5703–5710.

Marée, A.F.M., Grieneisen, V.A., and Edelstein-Keshet, L. (2012). How Cells Integrate Complex Stimuli: The Effect of Feedback from Phosphoinositides and Cell Shape on Cell Polarization and Motility. PLOS Computational Biology 8, e1002402.

McQuin, C., Goodman, A., Chernyshev, V., Kamentsky, L., Cimini, B.A., Karhohs, K.W., Doan, M., Ding, L., Rafelski, S.M., Thirstrup, D., et al. (2018). CellProfiler 3.0: Next-generation image processing for biology. PLOS Biology 16, e2005970.

Millius, A., and Weiner, O.D. (2009). Chemotaxis in neutrophil-like HL-60 cells. Methods Mol Biol 571, 167–177.

Moreau, H.D., Blanch-Mercader, C., Attia, R., Maurin, M., Alraies, Z., Sanséau, D., Malbec, O., Delgado, M.-G., Bousso, P., Joanny, J.-F., et al. (2019). Macropinocytosis Overcomes Directional Bias in Dendritic Cells Due to Hydraulic Resistance and Facilitates Space Exploration. Developmental Cell 49, 171–188.e5.

Niggli, V. (1999). Rho-kinase in human neutrophils: a role in signalling for myosin light chain phosphorylation and cell migration. FEBS Letters 445, 69–72.

Parent, C.A., and Devreotes, P.N. (1999). A cell’s sense of direction. Science 284, 765–770.

Prentice-Mott, H.V., Chang, C.-H., Mahadevan, L., Mitchison, T.J., Irimia, D., and Shah, J.V. (2013). Biased migration of confined neutrophil-like cells in asymmetric hydraulic environments. PNAS 110, 21006–21011.

Renkawitz, J., Kopf, A., Stopp, J., de Vries, I., Driscoll, M.K., Merrin, J., Hauschild, R., Welf, E.S., Danuser, G., Fiolka, R., et al. (2019). Nuclear positioning facilitates amoeboid migration along the path of least resistance. Nature 568, 546–550.

Simon, N., Friedman, J., Hastie, T., and Tibshirani, R. (2013). A Sparse-Group Lasso. Journal of Computational and Graphical Statistics 22, 231–245.

Smith, L.A., Aranda-Espinoza, H., Haun, J.B., Dembo, M., and Hammer, D.A. (2007). Neutrophil Traction Stresses are Concentrated in the Uropod during Migration. Biophys J 92, L58–L60.

Srinivasan, S., Wang, F., Glavas, S., Ott, A., Hofmann, F., Aktories, K., Kalman, D., and Bourne, H.R. (2003). Rac and Cdc42 play distinct roles in regulating PI(3,4,5)P3 and polarity during neutrophil chemotaxis. J Cell Biol 160, 375–385.

Swaney, K.F., Huang, C.-H., and Devreotes, P.N. (2010). Eukaryotic Chemotaxis: A Network of Signaling Pathways Controls Motility, Directional Sensing, and Polarity. Annu Rev Biophys 39, 265–289.

Tibshirani, R. (1996). Regression Shrinkage and Selection via the Lasso. Journal of the Royal Statistical Society. Series B (Methodological) 58, 267–288.

Tsai, T.Y.-C., Collins, S.R., Chan, C.K., Hadjitheodorou, A., Lam, P.-Y., Lou, S.S., Yang, H.W., Jorgensen, J., Ellett, F., Irimia, D., et al. (2019). Efficient Front-Rear Coupling in Neutrophil Chemotaxis by Dynamic Myosin II Localization. Dev. Cell 49, 189–205.e6.

Uchida, K.S.K., Kitanishi-Yumura, T., and Yumura, S. (2003). Myosin II contributes to the posterior contraction and the anterior extension during the retraction phase in migrating Dictyostelium cells. Journal of Cell Science 116, 51–60.

Van Keymeulen, A., Wong, K., Knight, Z.A., Govaerts, C., Hahn, K.M., Shokat, K.M., and Bourne, H.R. (2006). To stabilize neutrophil polarity, PIP3 and Cdc42 augment RhoA activity at the back as well as signals at the front. Journal of Cell Biology 174, 437–445.

Wang, F. (2009). The Signaling Mechanisms Underlying Cell Polarity and Chemotaxis. Cold Spring Harb Perspect Biol 1, a002980.

Wang, F., Herzmark, P., Weiner, O.D., Srinivasan, S., Servant, G., and Bourne, H.R. (2002). Lipid products of PI(3)Ks maintain persistent cell polarity and directed motility in neutrophils. Nat Cell Biol 4, 513–518.

Wang, Y., Ku, C.-J., Zhang, E.R., Artyukhin, A.B., Weiner, O.D., Wu, L.F., and Altschuler, S.J. (2013). Identifying network motifs that buffer front-to-back signaling in polarized neutrophils. Cell Rep 3, 1607–1616.

Weiner, O.D., Neilsen, P.O., Prestwich, G.D., Kirschner, M.W., Cantley, L.C., and Bourne, H.R. (2002). A PtdInsP(3)- and Rho GTPase-mediated positive feedback loop regulates neutrophil polarity. Nat Cell Biol 4, 509–513.

Weiner, O.D., Marganski, W.A., Wu, L.F., Altschuler, S.J., and Kirschner, M.W. (2007). An actin-based wave generator organizes cell motility. PLoS Biol 5, e221.

Xu, J., Wang, F., Van Keymeulen, A., Herzmark, P., Straight, A., Kelly, K., Takuwa, Y., Sugimoto, N., Mitchison, T., and Bourne, H.R. (2003). Divergent Signals and Cytoskeletal Assemblies Regulate Self-Organizing Polarity in Neutrophils. Cell 114, 201–214.

Yang, H.W., Collins, S.R., and Meyer, T. (2016). Locally excitable Cdc42 signals steer cells during chemotaxis. Nature Cell Biology 18, 191–201.

Zou, H., and Hastie, T. (2005). Regularization and variable selection via the elastic net. Journal of the Royal Statistical Society: Series B (Statistical Methodology) 67, 301–320.

